# Balancing grain yield trade-offs in ‘Miracle-Wheat’

**DOI:** 10.1101/2023.02.23.529729

**Authors:** Ragavendran Abbai, Guy Golan, C. Friedrich H. Longin, Thorsten Schnurbusch

**Author notes:** Correspondence: Ragavendran Abbai, Thorsten Schnurbusch.

## Abstract

Introducing variations in inflorescence architecture, such as the ‘Miracle-Wheat’ (*Triticum turgidum* convar. *compositum* (L.f.) Filat.) with a branching spike, has relevance for enhancing wheat grain yield. However, in the spike-branching genotypes, the increase in spikelet number is generally not translated into grain yield advantage because of reduced grains per spikelet and grain weight. Here, we investigated if such trade-offs might be a function of source-sink strength by using 385 RILs developed by intercrossing the spike-branching landrace TRI 984 and CIRNO C2008, an elite durum (*T. durum* L.) cultivar; they were genotyped using the 25K array. Various plant and spike architectural traits, including flag leaf, peduncle and spike senescence rate, were phenotyped under field conditions for two consecutive years. On Chr 5AL, we found a new modifier QTL for spike-branching, *branched head^t^ 3* (*bh^t^-A3*), which was epistatic to the previously known *bh^t^-A1* locus. Besides, *bh^t^-A3* was associated with more grains per spikelet and a delay in flag leaf senescence rate. Importantly, favourable alleles *viz., bh^t^-A3* and *grain protein content* (*gpc-B1*) that delayed senescence are required to improve grain number and grain weight in the spike-branching RILs. In summary, achieving a balanced source-sink relationship might minimise grain yield trade-offs in Miracle-Wheat.

**HIGHLIGHT:** Genetic interplay between sink number and post-anthesis source activity limits grain yield in the spike-branching ‘Miracle-Wheat’.

## INTRODUCTION

Wheat (*Triticum* sp.) inflorescence – ‘Spike’ is a determinate structure harbouring the grain-bearing spikelets on its rachis in a distichous pattern. During immature spike development, the inflorescence meristem gives rise to multiple spikelet meristems in an acropetal manner. In turn, each spikelet meristem (indeterminate) produces florets, that potentially form grains (Kirby and Appleyard, 1984; Koppolu and Schnurbusch, 2019). However, some exceptions deviate from this standard developmental programme, such as the ‘Miracle-Wheat’ that produces a non-canonical spike with lateral branches instead of spikelets. Here, due to a single amino acid substitution in the *branched head^t^* (*bh^t^*) allele of *T. turgidum* convar. *compositum* (L.f.) Filat. accessions, encoding an APETALA2/ETHYLENE RESPONSIVE FACTOR (AP2/ERF) transcription factor, the spikelet meristems lose their identity and determinacy while partially behaving as inflorescence meristems, producing lateral branches or multiple spikelets per rachis node (Poursarebani *et al*., 2015). Similarly, in hexaploid wheat, variations for supernumerary spikelet formation were also found for the wheat *FRIZZY PANICLE* (*WFZP*) (Dobrovolskaya *et al*., 2015), *Photoperiod-1 (Ppd-1)* (Boden *et al*., 2015), *TEOSINTE BRANCHED1 (TB1)* (Dixon *et al*., 2018), and *HOMEOBOX DOMAIN-2 (HB-2)* (Dixon *et al*., 2022). While branching spikes have considerably higher yield potential, i.e., more spikelet number, they often suffer from grain weight trade-offs, as observed in the tetraploid Miracle-Wheat (Poursarebani *et al*., 2015). Moreover, despite the increase in overall grain number per spike, spikelet fertility (grains per spikelet) decreased in response to spike-branching (Wolde *et al*., 2021).

A large body of evidence suggests that wheat grain yield is an outcome of multiple trait-trait interactions mediated by developmental, physiological and environmental factors across the entire lifespan, although some stages are more critical than others (Brinton and Uauy, 2019; Guo *et al*., 2017; Guo *et al*., 2018a; Guo *et al*., 2016; Guo *et al*., 2018b; Murchie *et al*., 2023; Reynolds *et al*., 2022; Slafer *et al*., 2023). They can broadly be classified as source and sink strength related, which jointly determine a particular genotype’s assimilate production and reallocation potential. Typically, green tissues of the plant – both foliar (leaves) and non-foliar (peduncle, spikes) are the photosynthesising organs that act as ‘source’ for resource generation (Chang *et al*., 2022; Molero and Reynolds, 2020). In the pre-anthesis phase, assimilates are partitioned to both vegetative biomass establishment and developing spikes – that determine the overall yield potential (Fischer, 2011; Slafer, 2003). The inflorescence architecture, *viz.,* spikelet number per spike, floret number per spikelet, carpel size, rachis length etc., are determined before anthesis (Brinton and Uauy, 2019; Kirby and Appleyard, 1984). For instance, the ovary size during flowering regulated floret and grain survival in a panel of 30 wheat genotypes (Guo *et al*., 2016). Likewise, the duration of leaf initiation, spikelet initiation and stem elongation period influenced spike fertility in bread wheat (Roychowdhury *et al*., 2023). The source strength is often characterised by radiation use efficiency (RUE), i.e., the ability for light interception and biomass production (Acreche and Slafer, 2009; Molero *et al*., 2019). However, the balance between the resources allocated to the ‘vegetative vs reproductive’ tissues largely dictates the yield potential (Dreccer *et al*., 2014; Ferrante *et al*., 2013), a trait that has been under selection throughout the history of wheat breeding. The deployment of semi-dwarf *Rht-1* alleles (‘green revolution’ gene) significantly increased the harvest index and the grain number per unit area, possibly by enhancing the flow of assimilates (as the stem length is considerably reduced) to the juvenile spikes (Fischer and Stockman, 1986; Slafer *et al*., 2023). However, other strategies might currently be required to further the resource allocation to early spike development as the semi-dwarf *Rht-1* allele is already a selection target (Peng *et al*., 1999). Increasing the harvest index in the genotypes with high biomass (more robust source) might enhance grain yield (Sierra-Gonzalez *et al*., 2021). Overall, the source strength from the terminal spikelet stage to anthesis majorly determines grain number and size in wheat.

After anthesis, the initiation of senescence in the foliar, but also non-foliar tissues drives extensive re-mobilization of resources into the developing grains; previous studies indicated that flag leaf and spike photosynthesis contribute to most of the assimilates during the grain filling phase (Distelfeld *et al*., 2014; Molero and Reynolds, 2020). Hence, a delayed senescence resulted in extended photosynthesis (functional stay-green), leading to higher grain yield (Chapman *et al*., 2021b; Christopher *et al*., 2016; Hassan *et al*., 2021; Kichey *et al*., 2007; Li *et al*., 2022; Niu *et al*., 2023; Yan *et al*., 2023). However, the effect of delayed senescence was not consistent; for instance, prolonged photosynthesis influenced grain yield attributes only under low nitrogen conditions (Derkx *et al*., 2012; Gaju *et al*., 2011). The *GPC-B1* locus encoding *NO APICAL MERISTEM (NAM)*, a *NAC* transcription factor that is the major regulator of senescence rate in wheat (Uauy *et al*., 2006); but, despite a 40% increase in flag leaf photosynthesis, the *NAM* RNAi wheat lines had no advantage in grain weight compared to the control plants (Borrill *et al*., 2015). In addition, the stay-green phenotype of *gpc-A1* and *gpc-D1* mutants did not influence grain yield determinants (Avni *et al*., 2014). However, (Chapman *et al*., 2021b) reported that novel *NAM-1* allele that delayed senescence was associated with 14% increase in the final grain weight, possibly by enhancing resource re-mobilization. A plausible explanation for such discrepancies might be that grain yield in wheat is largely sink-limited (Lichthardt *et al*., 2020; Reynolds *et al*., 2005); the surplus water-soluble carbons that remain in the stem at physiological maturity supports this hypothesis (Serrago *et al*., 2013). Thus, a higher sink capacity might be essential to capitalise on the extended photosynthetic period during the grain filling phase (Lichthardt *et al*., 2020). In this context, a reductionist approach that focusses on characterising individual component traits might assist in the deeper understanding of source-sink dynamics but also be integrated to pin-point favourable combinations of alleles/haplotypes for improving wheat grain yield (Brinton and Uauy, 2019; Reynolds *et al*., 2022).

As Miracle-Wheat has a stronger sink (more spikelet number), we hypothesized that delimited post-anthesis source strength might explain the spike-branching induced grain yield trade-offs. To examine this, we developed a bi-parental wheat population comprising about 385 RILs by crossing the spike-branching TRI 984 with an elite durum CIRNO C2008. The idea was to evaluate this population under field conditions for various architectural traits, as well as the senescence rate of the flag leaf, the peduncles and the spike (details are in ‘Materials and Methods’ section). In summary, our current study aims to explain: i. The relationship between senescence rate and trade-offs regulating grain yield (spike-branching–grain number–grain weight); ii. The underlying genetics of such trade-offs; iii. Favourable trait and allele combinations of relevant QTLs to balance grain yield trade-offs; and iv. Finally, to verify if spike-branching might be a potential selection target to enhance grain yield in wheat.

## MATERIALS AND METHODS

### Population development

A bi-parental population comprising 385 RILs was developed by crossing the spike-branching Miracle-Wheat accession, ‘TRI 984’ and elite durum from CIMMYT, ‘CIRNO C2008’ (hereafter referred to as ‘CIRNO’). A modified speed breeding method (Ghosh *et al*., 2018; Watson *et al*., 2018) was used for rapid generation advancement from F_3_ to F_5_. Initially, the grains were sown in the 96 well trays and grown in standard long day conditions *viz.,* 16h light (19^◦^C) and 8h dark (16^◦^C) for about two weeks. Later, the trays were transferred to speed breeding conditions *viz.,* 22h light (22^◦^C) and 2h dark (17^◦^C) to accelerate the growth. The spikes were harvested at maturity, and a similar method was used for the next cycle. Finally, the obtained F_5_ plants were multiplied under field conditions during the spring of 2020, and the resulting F_6_ grains (RILs) were genotyped and phenotyped.

### Genotyping and linkage map construction

The parental lines and three F_6_ grains per RIL were sown in 96 well trays and were grown in standard greenhouse conditions (16h light; 19^◦^C & 8h dark; 16^◦^C) for about two weeks. Leaves were sampled at the two-leaf stage from all the seedlings and stored at -80^◦^C until further use. During the sampling, the leaves from the three replications of a particular RIL were pooled, and genomic DNA was extracted. The DNA integrity was evaluated on agarose gel, after which about 50 ng/µl aliquots were prepared for the genotyping. Eventually, the 25K wheat SNP array from SGS-TraitGenetics GmbH was used for genotyping the 385 RILs along with the parental lines. However, only the 21K markers scored to the A & B sub-genome were considered for further analysis; we found that 5,089 makers were polymorphic (Fig. S1A). The linkage map was developed using the regression and maximum likelihood methods in JoinMap v4.1 (Stam, 1993). A subset of 2,128 markers was prepared after filtering, *viz.,* without segregation distortion (determined based on Chi-squared test), <10% heterozygosity and <10% missing (Fig. S1B&C). Haldane’s mapping function was used in the regression method, while the maximum likelihood method involved the spatial sampling thresholds of 0.1, 0.05, 0.03, 0.02 and 0.01 with three optimisation rounds per sample. Outcomes from both these methods were used to determine the 14 linkage groups and final map order.

### Examining plant and spike architectural traits

#### Greenhouse conditions

After collecting the leaves for genotyping (as described in the previous section), the two-week-old seedlings were vernalised at 4^◦^C for one month. Then, the seedlings were transferred to 9 cm square pots, grown in long day conditions (16h light; 19^◦^C & 8h dark; 16^◦^C). In both the parents – TRI 984 and CIRNO, tillers per plant (at booting) and spike number per plant (at maturity) were phenotyped. Besides, flag leaf verdancy was measured at eight different locations along the leaf blade (Borrill *et al*., 2019) at heading and also at 30 days after heading using the SPAD-502 chlorophyll meter (Konica Minolta). Standard fertilization was performed, and plants were treated with pesticides based on the requirement.

#### Field conditions

The genotypes were screened at IPK-Gatersleben (51^°^49ʹ23ʹʹN, 11^°^17ʹ13ʹʹE, 112m altitude) under field conditions for two growing seasons *viz.,* the F_6_ derived RILs in the spring of 2021, and the F_7_ derived RILs in the spring of 2022. They were grown in an α-lattice design with three replications, while each 1.5 m^2^ plot had six 20 cm spaced rows comprising two genotypes (three rows each). Standard agronomic and management practices were in place throughout the growth cycle; however, the experimental trial was completely rainfed. Besides, a subset of genotypes (F_6_ derived RILs) in one replication was evaluated at the University of Hohenheim (48^°^42’50’’N, 9^°^12’58’’E, 400 m altitude) in 2022.

Plants from the inner rows (at least five measurements per plot) were considered for all the phenotyping except for grain yield per meter row, where the mean of all three rows of a particular genotype was measured. Days to heading (DTH) was determined at ‘Zadoks 55’, i.e. when half of the spike has emerged (Zadoks *et al*., 1974) in about 50% of the plants in a particular plot. Later, this was converted into growing degree days (GDD) to account for temperature gradients (Miller *et al*., 2001). The distance from the tip of the flag leaf to its base was considered as the flag leaf length, while the flag leaf width was the end-to-end horizontal distance at the middle of the leaf. At heading, flag leaf verdancy was measured using the SPAD-502 chlorophyll meter (Konica Minolta) at eight different locations along the leaf blade (Borrill *et al*., 2019). Flag leaf senescence was screened at 30 days after heading using a four-point severity scale from ‘1’ indicating the least senescence to ‘4’ for the highest senescence (Fig. S2A). The number of senesced peduncles per 10 peduncles was counted from the inner rows to determine peduncle senescence (%) (Chapman *et al*., 2021a). In this context, we found a gradient of yellowness in the peduncle across the RILs; however, in the current study, this was not differentiated, i.e. we had only two classes – green and yellow (Fig. S2B). Days to maturity (DTM) was determined when most spikes turned yellow in a particular plot; later, this was converted to growing degree days similar to days to heading.

Spike weight, spike length (without awns) and straw biomass (dry weight of culm along with leaves) were measured after harvest. In addition, a scoring method was developed for estimating supernumerary spikelets (two spikelets per rachis node) and spike-branching (true branching with mini-spikes from the rachis nodes) (Fig. S3). ‘0’ (standard spike), ‘1’ (supernumerary spikelets only at the basal part of the spike), ‘2’ (supernumerary spikelets until half of the spike), ‘3’ (supernumerary spikelets throughout the spike) and ‘4’ (proper branching). Floret number was measured from the non-branching genotypes from two spikelets at the centre of the spike at harvest. Besides, derived traits such as grains per spikelet, grain filling duration (Chapman *et al*., 2021a), and harvest index was calculated as follows:

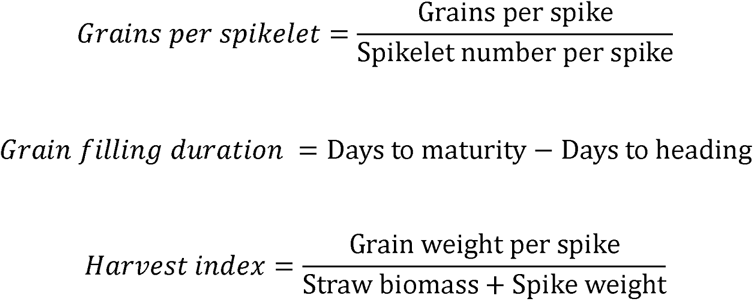

’Marvin’ digital grain analyser (GTA Sensorik GmBH, Neubrandenburg, Germany) was used to determine grains per spike, thousand-grain weight, grain length, and grain width. We also recorded the grain width and length of the parental lines manually using a Vernier calliper to reconfirm the observed trend from the ‘Marvin’ digital seed analyser (Fig. S4). All the above-mentioned traits were recorded at IPK-Gatersleben, while only the spike architectural traits were phenotyped from the experiment conducted at the University of Hohenheim.

### Phenotypic and Genetic analyses

Genstat 19 (VSN International, Hemel Hempstead, UK) and GraphPad Prism 9.3.1 (GraphPad Software, San Diego, California, USA) were used for all the statistical analyses. Ordinary one-way ANOVA followed by Dunnett’s multiple comparisons test was employed for multiple-range comparisons, whereas an unpaired *Student’s t-test* was used to compare two groups. Pearson correlation was used to study the relationship among the traits of interest; besides, simple linear regression assisted in understanding the effect of a particular trait (explanatory variable) on another (response variable). The corresponding figures contain all the relevant details, such as P-value, R^2^, and the number of samples compared.

QTL mapping was performed in Genstat 19 using the following criteria: i. step size of 10 cM, ii. minimum cofactor proximity of 50 cM, iii. minimum QTL separation distance of 30 cM and iv. genome wide significance ‘α =0.05’. Simple interval mapping (SIM) was performed as an initial scan to determine the positions of potential candidate QTL(s). These positions were used as cofactors for multiple rounds of composite interval mapping (CIM); CIM was repeated until similar results were obtained at least three consecutive times. Finally, QTL backward-selection was carried out after CIM to estimate various QTL effects, including the determination of QTL interval, high-value allele, additive effects, and phenotypic variance explained. The QTLs were then visualised using MapChart 2.32 (Voorrips, 2002).

## RESULTS

### Spike-branching affects grains per spikelet and thousand-grain weight

Consistent with previous findings using different germplasm (Wolde *et al*., 2021), while the spike-branching landrace TRI 984 had more spikelets and florets per spike, the spikelets contained fewer florets and grains than the elite durum cultivar CIRNO (Fig. 1A-D and Fig. S5A-C). However, we found no difference in grain number per five spikes, but a considerably reduced thousand-grain weight associated with shorter grains was observed in TRI 984 (Fig. 1E-H). While CIRNO flowered earlier (Fig. 1I), it had greener flag leaves both at heading (Fig. 1J) and also after 30 days of heading (Fig. 1K) along with greener peduncles (Fig. 1L)’. Besides, CIRNO had longer but narrower flag leaves (Fig. S5D&E), fewer tillers (Fig. 1M), yet spikes per plant remained unaltered (Fig. 1N), shorter spikes (Fig. S5F) and shorter plant stature (Fig. S5G) as opposed to TRI 984. We found considerable differences in spike-branching expressivity in TRI 984 (Fig. S5H), which can explain the grain number, grain weight and spike weight variations observed among various replications of TRI 984 (Fig. 1E-H, O&P). Such inconsistencies in spike-branching has also been reported earlier in another Miracle-Wheat – Elite cultivar biparental population (Wolde et al., 2021). Furthermore, there was no difference in the average spike weight (n=5) (Fig. 1O), but CIRNO had more grain yield per five spikes (Fig. 1P). These observations indicate a clear difference in terms of assimilate production and reallocation patterns between the two genotypes. Variations in tiller number (at booting) might indicate different resource partitioning strategies in TRI 984 and CIRNO during the pre-anthesis phase; however, there was no difference in the total number of spikes at maturity. Importantly, the spike-branching landrace TRI 984 exhibited a shorter grain filling period (quicker senescence), which implies reduced resource production and reallocation compared to CIRNO (delayed senescence) during the post-anthesis phase. Besides, the resources required to maintain the vegetative parts might be higher in the case of TRI 984 because of the taller plant architecture than CIRNO. Based on these observations, it is conceivable that genetic analysis of the corresponding landrace-elite recombinants (TRI 984 x CIRNO) that vary in source-sink balance might provide mechanistic insights into the negative effect of spike-branching on grain number per spikelet and grain weight, two major components of final grain yield.

**Fig. 1.**
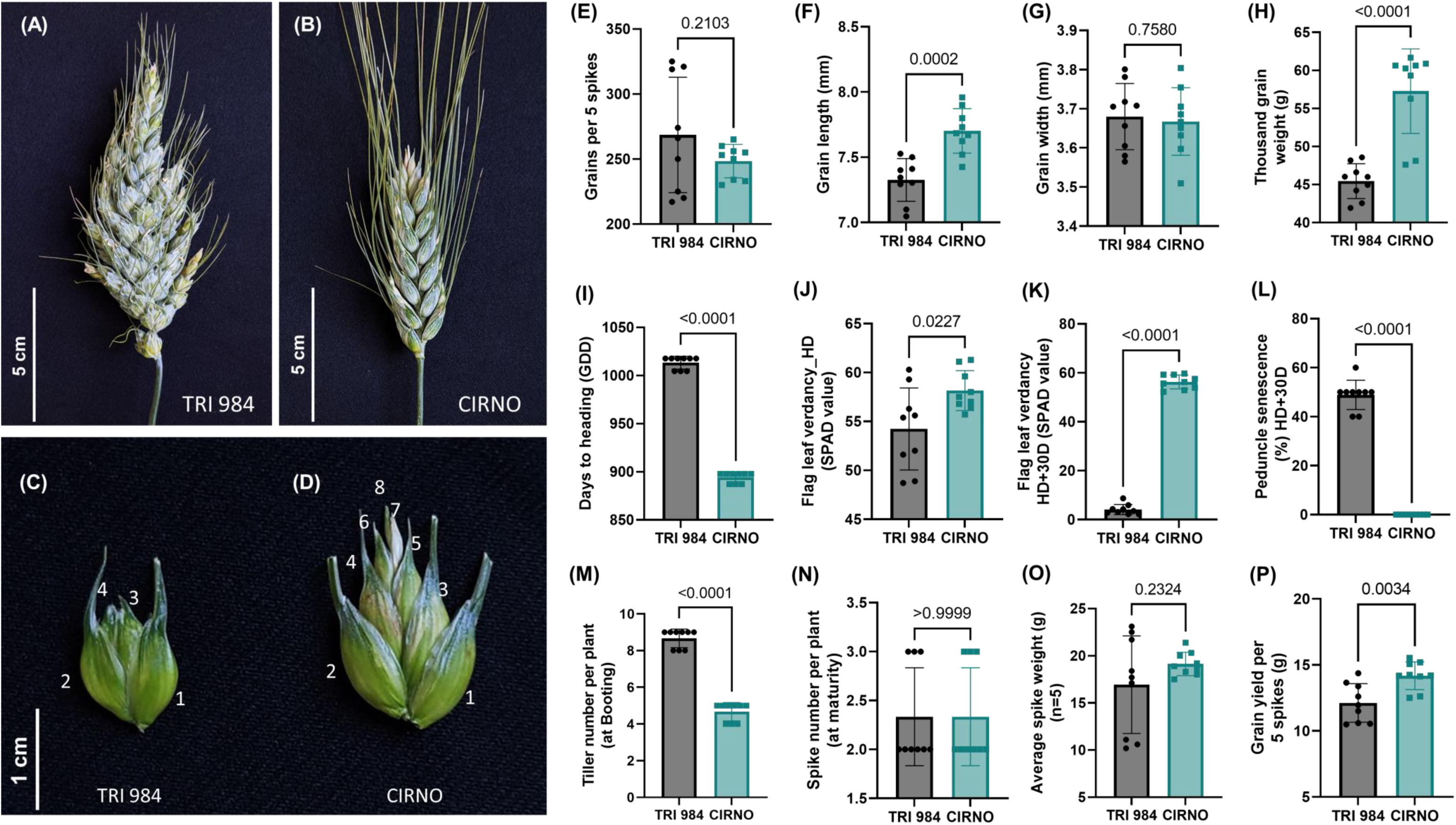
TRI 984 has a poor source-sink balance as opposed to CIRNO. (A) Spike-branching phenotype of TRI 984 and (B) Standard spike of CIRNO. (C) Spikelet from the central part of the TRI 984 main rachis shows reduced florets relative to (D) a spikelet from a similar position in CIRNO. (E) Despite having more spikelets, there is no increase in grain number per 5 spikes in TRI 984. (F-H) However, the grains are smaller in TRI 984, leading to a reduction in thousand-grain weight. (I) TRI 984 exhibited delayed heading, (J) lower flag leaf verdancy at heading, (K, L) accelerated flag leaf and peduncle senescence. TRI 984 had (M) more tillers at booting, (N) but final spike number per plant remained unaltered, (O) There was no difference in spike weight. However, (P) grain yield per five spikes was reduced in TRI 984 compared to CIRNO. Note: unpaired t-test was performed to determine significance in (E-P) and the resulting P-values (Two-tailed analysis) are displayed. Data represented in (J, K, M, & N) were obtained from green-house experiment, while the all remaining traits were phenotyped from field grown plants.

### Grains per spikelet and thousand-grain weight are associated with senescence rate

As expected, we witnessed a considerable diversity for all the plant and spike architectural traits (Fig. S6). Importantly, flag leaf and peduncle senescence rates were independent of the heading date; this implies that there is a possibility for the lines that flowered late to senesce early and vice-versa (Fig. 2A). The lines with delayed flag leaf senescence also had the tendency of retaining green/verdant peduncles for a longer duration (Fig. 2A). In addition, the intensity of flag leaf greening (SPAD meter value) at heading had no effect (R^2^=0.012; p=0.212) on the progress of senescence (scored at 30 days after heading), indicating that these traits are largely independent (Fig. 2B). Flag leaf length and delay in senescence were positively related (R^2^=0.045; p=0.0034), while flag leaf width did not influence the same (Fig. S7A&B). Moreover, we observed that the lines with more verdant/greener flag leaves at heading (higher SPAD value) also had a more significant number of florets per spikelet (R^2^=0.085; p=0.0014), in line with the expected consequence of source strength on sink organ establishment before anthesis (Fig. S7C). Intriguingly, the number of florets and grains per spikelet, which is determined earlier, was somewhat associated with senescence rate, i.e., the lines with more florets and grains per spikelet tended to display delayed flag leaf senescence (R^2^=0.071; p=0.0007 & R^2^=0.14; p<0.0001) (Figs. S7D & 2C). Likewise, it has been previously reported that higher grain number increased the post-anthesis radiation use efficiency in wheat (Reynolds *et al*., 2005). Here, we mapped a QTL on Chr 5A (*bh^t^-A3*) influencing grains per spikelet and flag leaf senescence rate (Table S1). This posssibly implies a gene/QTL mediated pleiotropic association between sink number and flag leaf senescence rate, although these traits might not be physiologically dependent. Besides, the delayed senescence rate had a positive effect on thousand-grain weight (R^2^=0.13; p<0.0001) (Fig. 2D). We realised that the observed increase in thousand-grain weight is primarily due to the change in grain width (R^2^=0.08; p<0.0001) (Fig. 2E) and not grain length (Fig. S7E), suggesting that grain width is more plastic, influenced by resource reallocation compared to grain length. Nevertheless, it is clear that the longer duration of green flag leaf and peduncle is not simply ‘cosmetic’ – it influences grain yield determinants. This vital evidence supports our hypothesis that dissecting the source-sink relationship might have relevance in balancing the trade-offs that negatively regulate the final grain yield in ‘Miracle-Wheat’ like genotypes.

**Fig. 2.**
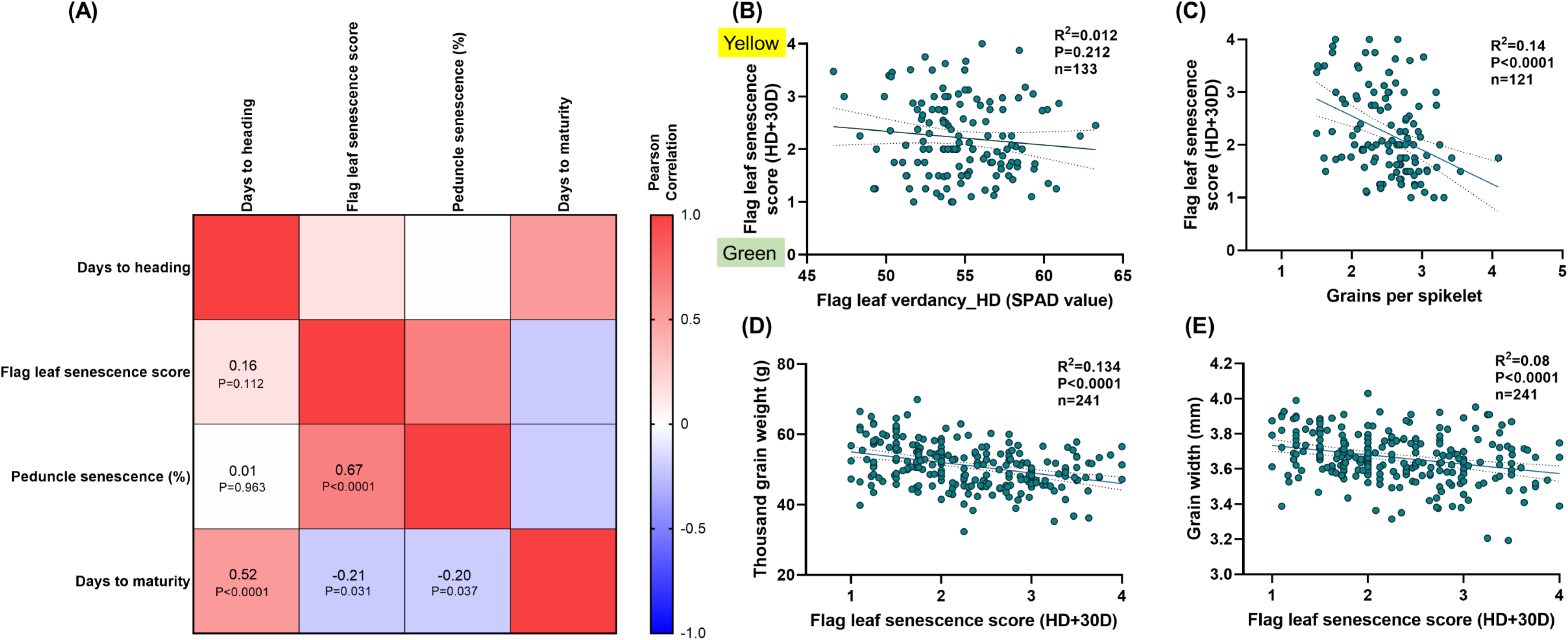
Functional ‘Stay-green’ phenotype was observed in the landrace-elite recombinants. (A) Flag leaf and peduncle senescence were positively related, independent of days to heading. (B) Flag leaf verdancy at the heading did not impact the progression of senescence. (C) Grains per spikelet was associated with flag leaf senescence rate and (D) the RILs with delayed senescence had higher thousand-grain weight, due to (E) wider grains. Note: (B-E) are linear regression plots with the explanatory variable on the x-axis, while the y-axis represents the response variable. R^2^ is the phenotypic variance explained, and the corresponding P-values of the regression analysis are displayed. The data presented in (A-E) were phenotyped from field grown plants.

#### The bh^t^-A1 locus underlies sink and source capacity

Using a scoring method based on the phenotype of the spike (Fig. S3), we mapped a major effect QTL for spike-branching on Chr 2A (Fig. 3A, Table S1) associated with the previously known locus *bh^t^-A1* (Poursarebani *et al*., 2015). Regardless of the increase in spikelet number per spike owing to the lateral branching (Fig. 3B), there was no difference in the total grains per spike (Fig. 3C); but, the *bh^t^-A1* locus was associated with a reduction in grain length (Fig. 3D) and thousand-grain weight (Fig. 3E). While the *bh-A1* lines had slightly longer flag leaf blades (Fig. 3F), the flag leaf verdancy at heading (Fig. 3G) and spike length (Fig. 3H) were negatively affected. Thus, the TRI 984 allele induced spike-branching and in addition might also affect the source capacity. Eventually, there was no advantage in the final grain yield because of spike-branching.

**Fig. 3.**
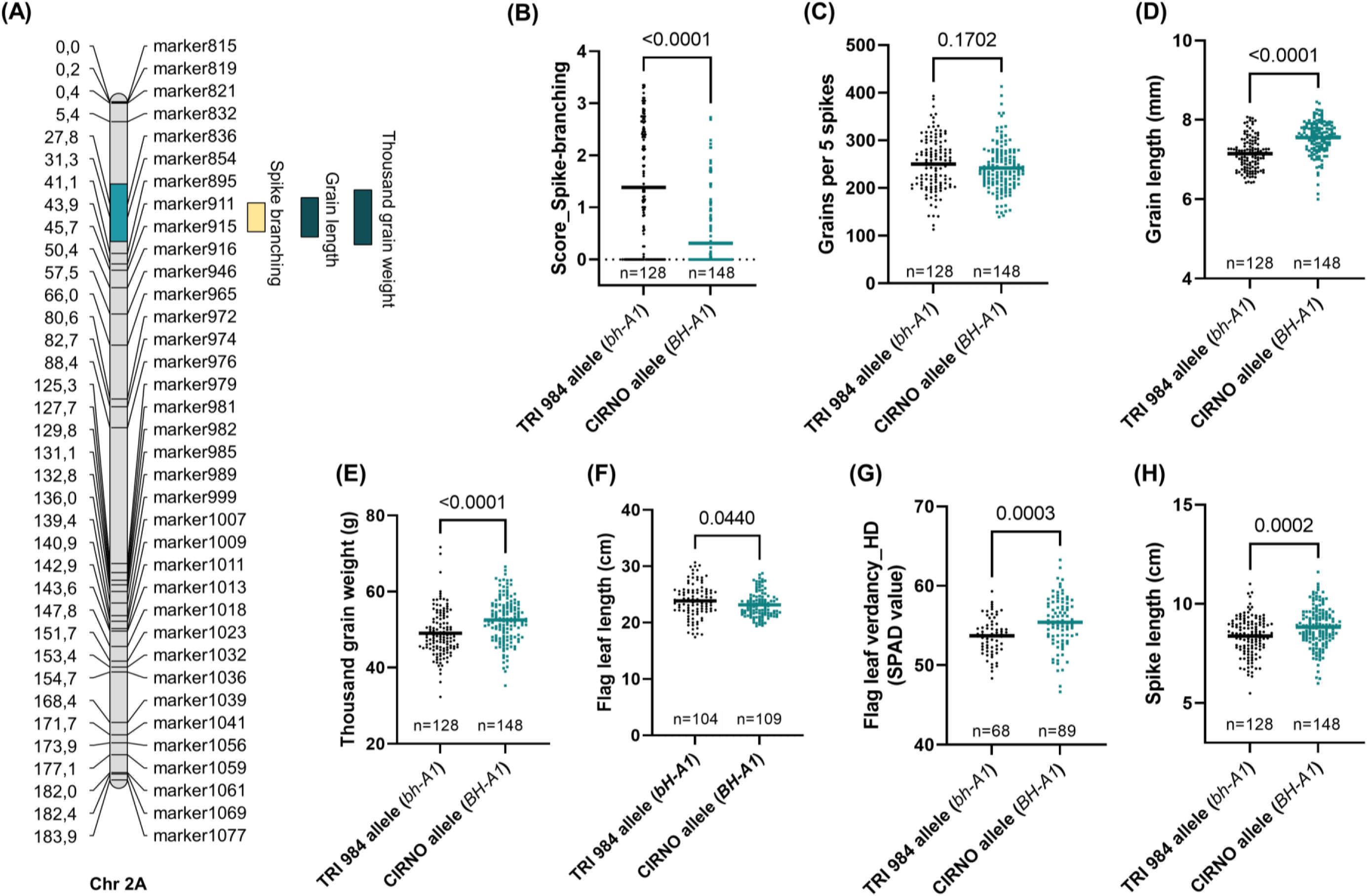
*bh^t^-A1* induces spike-branching but with a grain weight trade-off. (A) A major effect QTL hotspot for spike-branching, grain length and weight was mapped on the short arm of Chr 2A. (B) RILs with the TRI 984 allele showed spike-branching, (C) no difference in grains per 5 spikes, but a reduction in (D) grain length, (E) thousand-grain weight, (F) flag leaf length, (G) flag leaf verdancy at heading and (H) spike length. Note: In (B-H), ‘n’ represents the number of RILs that were compared for each allele class, *viz.,* TRI 984 allele data points are in ‘black’, while ‘cyan’ colored data points represent CIRNO allele. ‘Unpaired t test’ was used to determine the statistical significance, and the resulting P-values (Two-tailed analysis) are displayed for all the comparisons. Data obtained from field grown plants was used for the analysis in (A-H).

#### bh^t^-A3, a novel spike-branching locus on Chr 5A reshapes source-sink dynamics

Following similar phenotyping (Fig. S3), we mapped a QTL for spike-branching on the long arm of Chr 5A; here, the CIRNO allele contributed to the spike-branching phenotype (Fig. 4A, Table S1). We named the newly identified spike-branching modifier locus as ‘*bh^t^-A3*’ following the previously known *bh^t^-A1* (Poursarebani *et al*., 2015) and *bh^t^-A2* (Wolde et al., 2021) loci. Interestingly, the spike-branching effect of the *bh^t^-A3* locus manifests only in the presence of the mutated *bh^t^-A1* allele (Fig. 4A; Fig. S8A-D; Table S1). We divided the RILs into two sub-groups for QTL mapping *viz.,* by fixing i. *bh^t^-A1,* ii. *BH^t^-A1* and the outcome confirmed the epistasis of the *bh^t^-A3* to *bh^t^-A1* locus (Fig. S8A). Possibly, this indicates that the plasticity for spike-branching is introduced by *bh^t^-A1*, i.e., it might be first essential to have *bh^t^-A1* to disrupt the spikelet meristem identity and only then the *bh^t^-A3* locus might modify the branching intensity in the spikes. Besides, the grain number increase is only observed in the spike-branching RILs – when *bh^t^-A1* is present (Fig. S8C). Moreover, in this region, we found co-localised QTLs for an array of traits influencing source-sink dynamics. The CIRNO allele was associated with spike-branching (Fig. 4B), delayed flag leaf senescence (Fig. 4C), more extended grain filling period (Fig. 4D), increased grains per spikelet (Fig. 4E) and grain yield per five spikes (Fig. 4F). Besides, we also found a subtle, yet positive effect on grain width (Fig. S9A), florets per spikelet (Fig. S9B), straw biomass (Fig. S9C) and harvest index (Fig. S9D); but, the days to heading was not affected (Fig. S9E). Interestingly, we found that the observed variations in flag leaf senescence, grain filling duration and grain width were not dependent on the presence of *bh^t^-A1* (Fig. S8E-G). This pattern implies that the phenotypic variation explained by the 5A QTL hotspot for spike-branching expressivity and senescence rate might be the outcome of at least two linked genes. Taken together, this trend suggests that the favourable CIRNO allele (*bh^t^-A3*) mediates enhanced assimilate production and reallocation of the resources to sink organs, including the lateral branches/supernumerary spikelets because of longer grain filling duration.

**Fig. 4.**
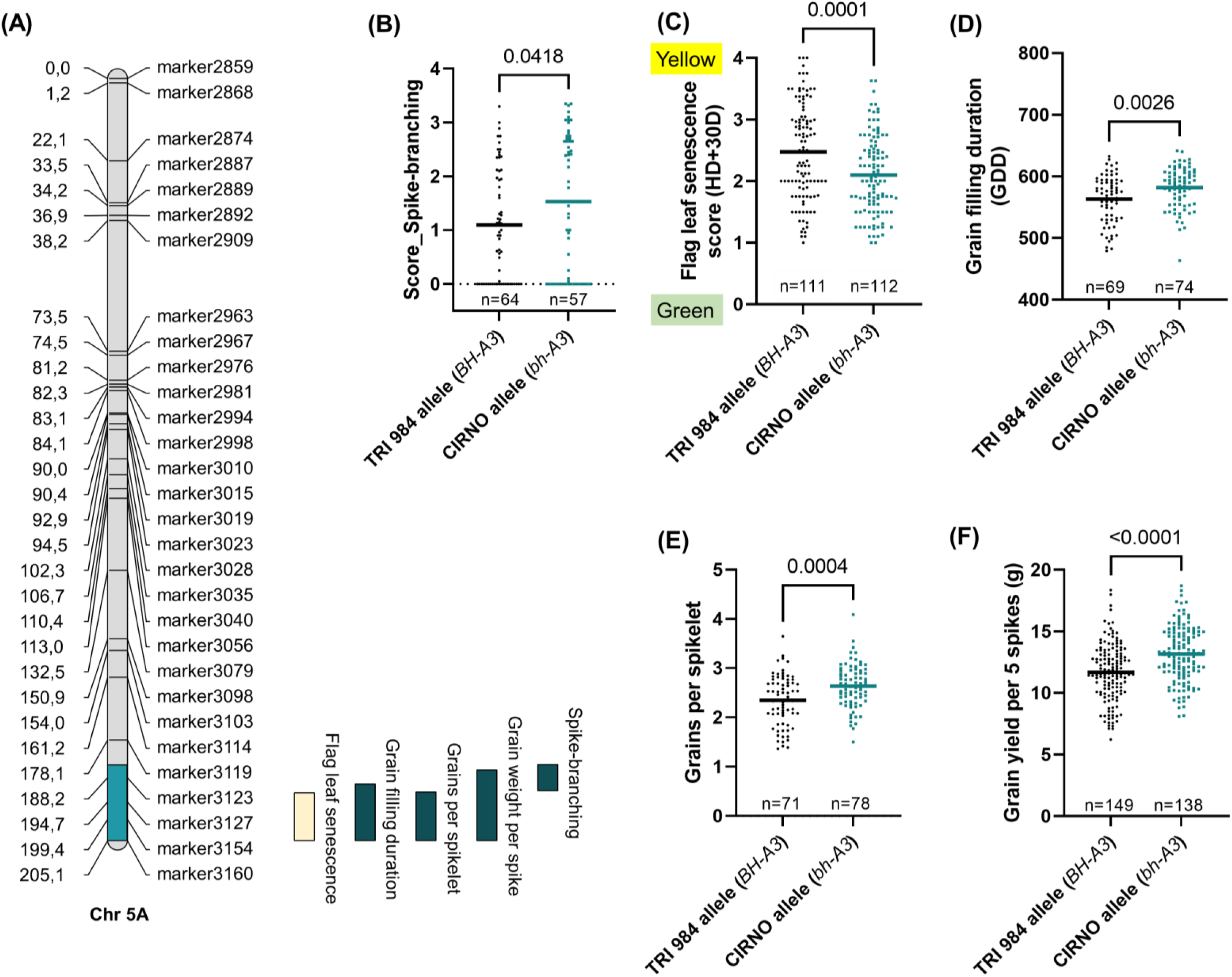
*bh^t^-A3*, a new modifier locus for spike-branching. (A) *bh^t^-A3* mediates spike-branching, flag leaf senescence rate, grain filling duration, grains per spikelet and grain yield per spike. The CIRNO allele (B) increases the expressivity of spike-branching (when *bh^t^-A1* is present), (C) delays flag leaf senescence rate, (D) increases grain filling duration, (E) grains per spikelet, and (F) grain yield per five spikes. Note: In (B-F), ‘n’ represents the number of RILs that were compared for each allele class *viz.,* TRI 984 allele data points are in ‘black’, while ‘cyan’ colored data points represent CIRNO allele. ‘Unpaired t test’ was used to determine the statistical significance, and the resulting P-values (Two-tailed analysis) are displayed for all the comparisons. Data obtained from field grown plants was used for the analysis in (A-F).

#### GPC-B1 is the major determinant of senescence rate and thousand-grain weight

A QTL on Chr 6B, which most likely is associated with *GPC-B1* (Uauy *et al*., 2006), explained most of the observed phenotypic variance for the overall plant senescence rate (Fig. 5A; Table S1). Likewise, it was found that mutations in the NAC domain of *NAM-A1* (*GPC-A1*) delayed peduncle and flag leaf senescence (Harrington *et al*., 2019). In the current study, the CIRNO allele ensured delay in the flag leaf (Fig. 5B), peduncle (Fig. 5C) and spike senescence (days to maturity) (Fig. 5D). Therefore, there might be a possibility of more reallocation into the sink organs, leading to an increase in grain width (Fig. 5E) and grain length (Fig. 5F). Accordingly, we observed a considerably higher thousand-grain weight in the RILs that senesce late (Fig. 5G). Besides, there was no meaningful difference in grain number per five spikes (Fig. S10A), straw biomass (Fig. S10B) and harvest index (Fig. S10C). Although flag leaf length was not directly affected by this QTL (Fig. S10D), longer flag leaves, in general, had a positive effect on thousand-grain weight in both allele groups (Fig. S11A&B) *viz., GPC-B1* (R^2^=0.11; p=0.0003) and *gpc-B1* (R^2^=0.043; p=0.041). Notably, the relationship between flag leaf length and grain weight was relatively stronger in the early senescing genotypes. In fact, the contribution to thousand-grain weight per unit length of flag leaf was higher in the case of *gpc-B1* allele that is associated with delayed senescence (Fig. S11C). Furthermore, we also included the effect of the allelic status at *bh^t^-A1*, another major QTL for grain weight, in addition to *gpc-B1*. Here, the RILs with various allele combinations revealed a similar positive relationship between flag leaf length and grain weight (Fig. 11D-G) *viz., BH^t^-A1*+*gpc-B1* (R^2^=0.113; p=0.0118), *BH^t^-A1*+*GPC-B1* (R^2^=0.164; p=0.0032), *bh^t^-A1*+*gpc-B1* (R^2^=0.12; p=0.0586), and *bh^t^-A1*+*GPC-B1* (R^2^=0.136; p=0.0048). Overall, these observations support the idea for the importance of source strength i.e., possibly more resource production and reallocation (delayed senescence) enhanced *per se* thousand-grain weight, including the spike-branching genotypes evaluated in the current population.

**Fig. 5.**
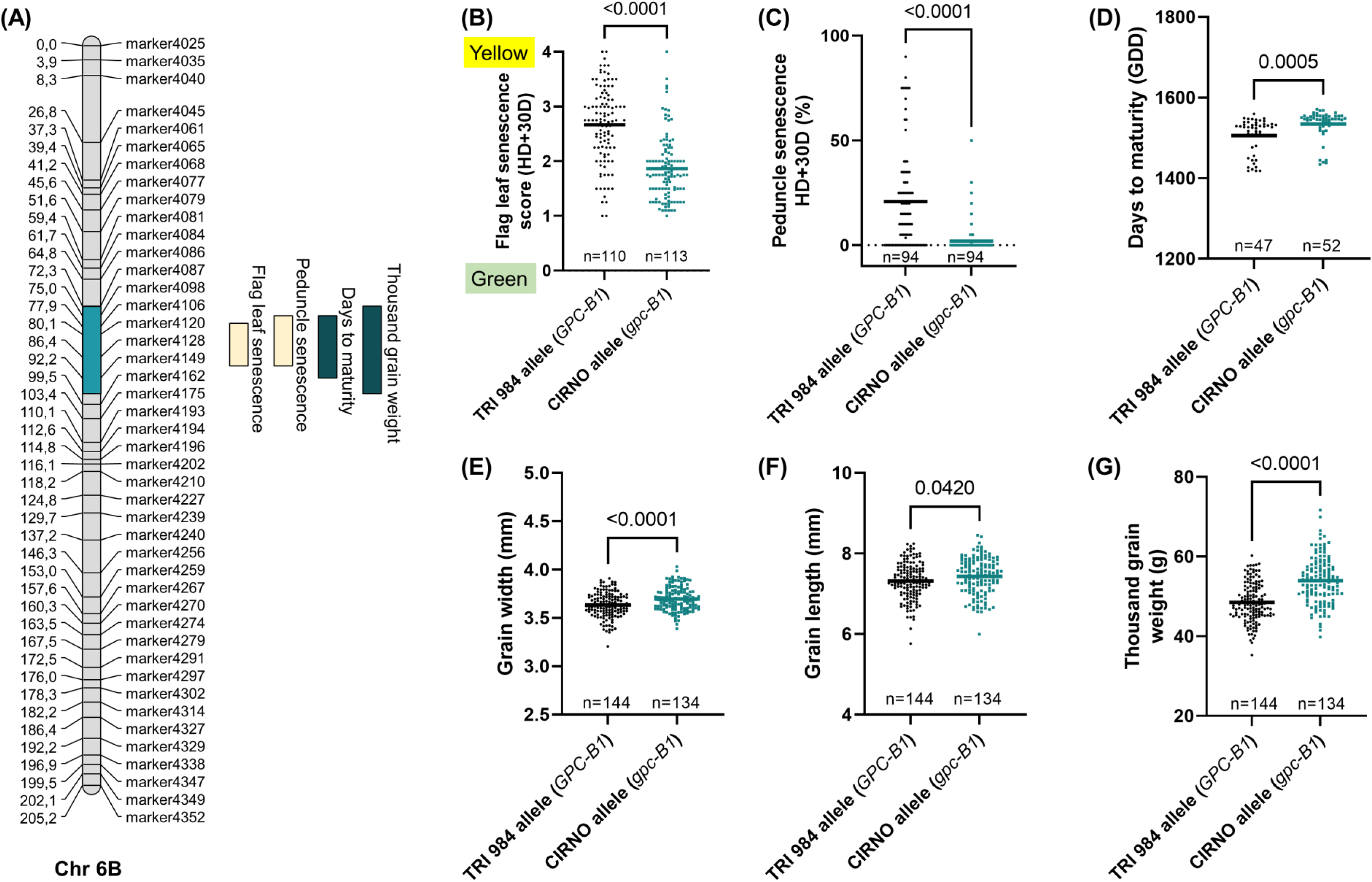
*gpc-B1* regulates senescence rate and grain weight. (A) QTLs for overall senescence rate and thousand-grain weight co-localised on Chr 6B. (B) The modern (CIRNO) allele mediated delay in flag leaf senescence, (C) peduncle senescence and (D) days to maturity (spike senescence). The resulting increase in the post-anthesis phase is translated into (E) an increase in grain width, (F) grain length and eventually (G) thousand-grain weight. Note: In (B-G), ‘n’ represents the number of RILs that were compared for each allele class, *viz.,* TRI 984 allele data points are in ‘black’, while ‘cyan’ colored data points represent CIRNO allele. ‘Unpaired t test’ was used to determine the statistical significance, and the resulting P-values (Two-tailed analysis) are displayed for all the comparisons. Data obtained from field grown plants was used for the analysis in (A-G).

#### Specific additive and epistatic interactions may increase yield potential in spike-branching genotypes

As the QTLs on Chr 2A, 5A and 6B explain variations in key source-sink attributes, we analyzed their various allelic combinations to better understand the grain yield trade-offs in spike-branching genotypes (Figs. 6A-H, 7A-G & 8). Interestingly, the spike-branching lines carrying *bh^t^-A1* and *bh^t^-A3* loci along with *gpc-B1* had higher grain number per five spikes (Fig. 6A, E) and were associated with a delay in post-anthesis flag leaf senescence. However, the difference in thousand-grain weight was observed only at IPK (Figs. 6B & S12A, B), while this effect was absent in Hohenheim (Figs. 6F & S12C, D). Nevertheless, they had higher grain yield per five spikes (Fig. 6C, G) across all the three environments *viz.,* IPK-2021, IPK-2022 and University of Hohenheim-2022 as opposed to the early senescing branched spike RILs (*bh^t^-A1+BH^t^-A3+GPC-B1*). Moreover, grain yield (per meter row) was also higher in the stay-green spike-branching RILs than the ones that senesced early (Fig. 6D).

**Fig. 6.**
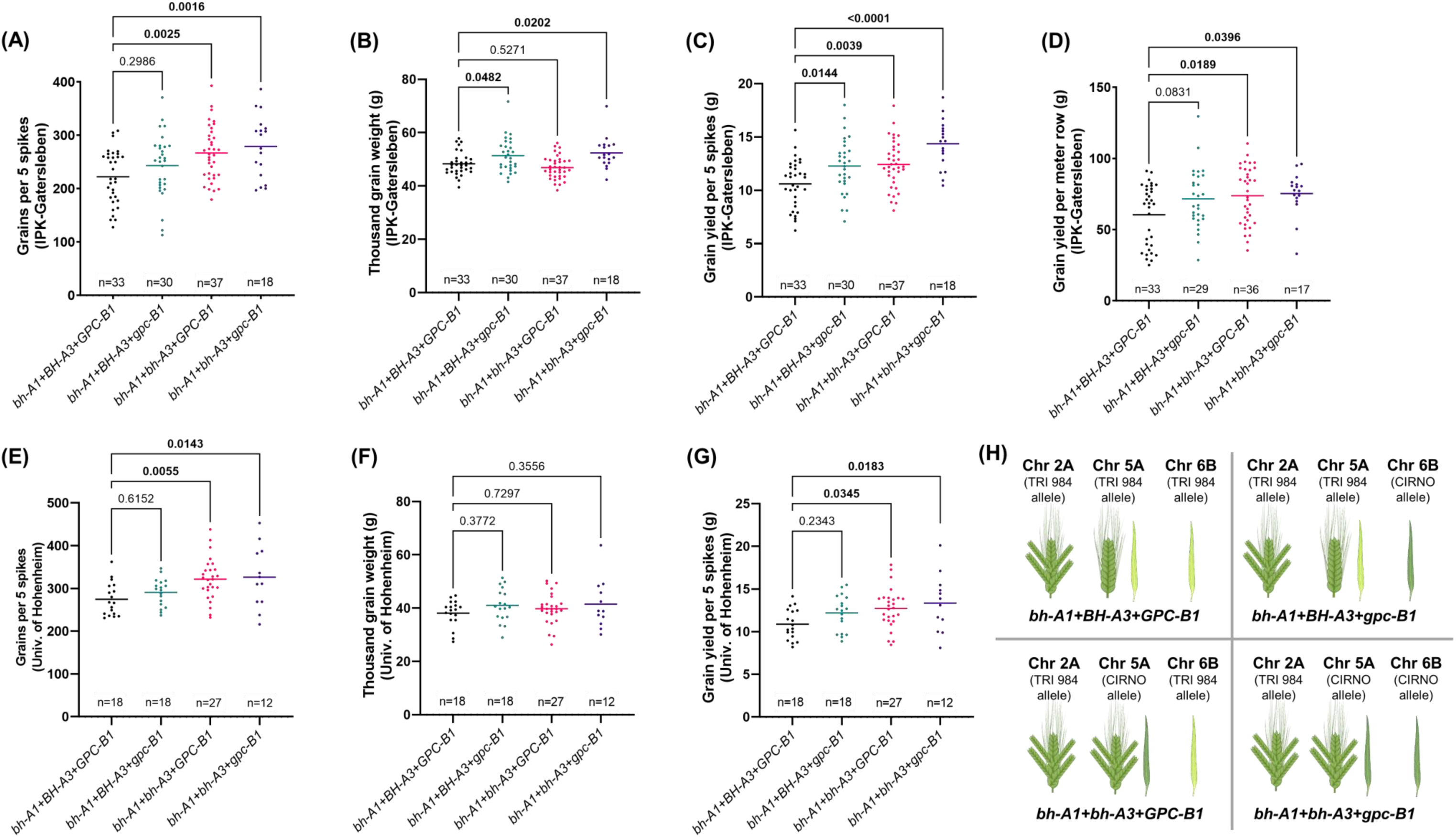
*bh^t^-A1*, *bh^t^-A3*, and *gpc-B1* balance the grain yield trade-offs in spike-branching recombinants. The RILs with various combinations of alleles were phenotyped at IPK-Gatersleben (2021 & 2022) and the University of Hohenheim (2022). At IPK, the spike-branching RILs that senesce late *(bh^t^-A1+bh^t^-A3+gpc-B1)* had (A) higher grain number per 5 spikes; (B) increased thousand-grain weight at IPK, (C) more grain yield per five spikes and (D) Finally, grain yield per meter row was also higher in the stay-green spike-branching RILs (calculated only at IPK). Likewise, at Hohenheim, (E) we observed more grains per five spikes, (F) but no change in thousand-grain weight; eventually, (G) there was an increase in grain yield per spike. (H) pictorial depiction of the various allelic combinations that are analysed in (A-G). Note: In (A-G), one-way ANOVA followed by Dunnett’s test was used to determine the statistical significance. All the comparisons are made with respect to ‘*bh^t^-A1+BH^t^-A3+GPC-B1’* allelic combination, and the corresponding P-values are displayed in all the graphs (significant ones are in bold). Data obtained from field grown plants was used for the analysis in (A-G). The image (H) is partly created using biorender (https://biorender.com/). *bh^t^-A1+BH^t^-A3+GPC-B1*: Recombinants with one locus for spike-branching and two loci for accelerated senescence; *bh^t^-A1+BH^t^-A3+gpc-B1*: Recombinants with one locus each for spike-branching and delayed senescence; *bh^t^-A1+bh^t^-A3+GPC-B1*: Recombinants with two loci for spike-branching, while carrying one locus for delayed senescence; *bh^t^-A1+bh^t^-A3+gpc-B1*: Recombinants with two loci each for spike-branching and delayed senescence.

Next, we compared the impact of delayed senescence between the spike-branching RILs and those with standard spikes (no spike-branching). Although, the spike-branching RILs carrying the favourable alleles (’*bh^t^-A1*+*BH^t^-A3*+*GPC-B1*’) had higher grain number (Fig. 7A), they had reduced thousand grain weight (Fig. 7B) than the lines with standard spikes having similar senescence rate i.e., ‘*BH^t^-A1*+*bh^t^-A3*+*gpc-B1*’; nevertheless, the final spike grain yield (Fig. 7C) was similar in both these cases. Finally, we compared the performance of these spike-branching RILs with both the parental lines. Here, we observed a non-significant increase in grain number, but a slightly reduced thousand grain weight than CIRNO (Fig. 7D, E). However, there was an increase in thousand grain weight and no change in grain number when compared to TRI 984 (Fig. 7D, E). Eventually, the grain yield per spike of the favourable spike-branching RILs was higher than TRI 984 and similar to CIRNO (Fig. 7F).

**Fig. 7.**
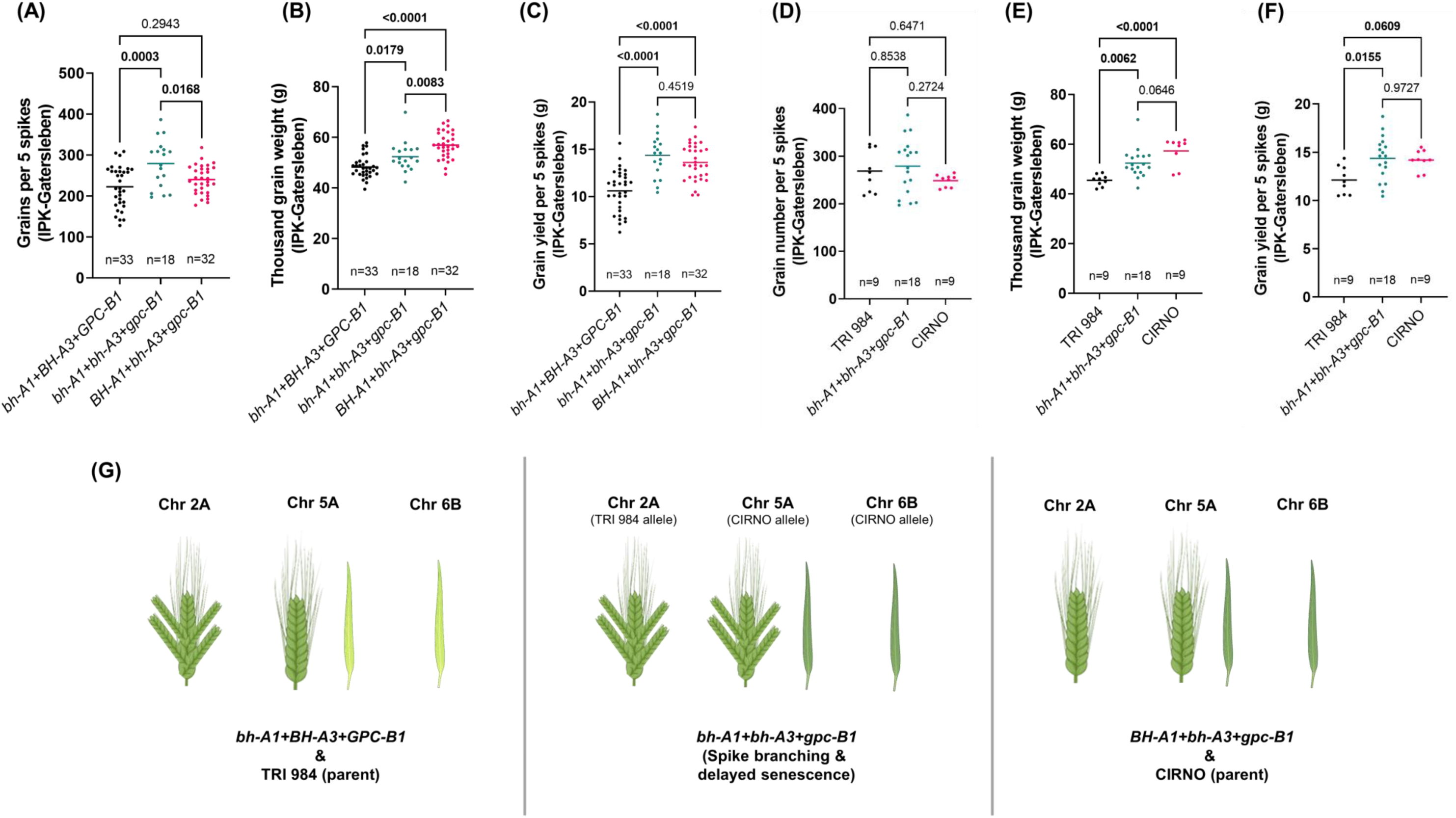
The favourable spike-branching RILs (*bh^t^-A1+bh^t^-A3+gpc-B1*) performed better than TRI 984, but similar to CIRNO. (A) The spike-branching RILs with delayed senescence produced more grains per spike, (B) but with a reduction in thousand grain weight than the corresponding genotypes with standard spike i.e., no spike-branching (*BH^t^-A1+bh^t^-A3+gpc-B1*). (C) Further, the spike grain yield was similar among the genotypes that senesce late, irrespective of the presence or absence of spike-branching. (D) However, there was no difference in grain number as opposed to TRI 984 and CIRNO. (E) The favourable spike-branching genotypes had higher thousand grain weight than TRI 984, whereas a reduction compared to CIRNO. (F) Finally, the spike grain yield was higher than TRI 984, but similar to CIRNO. (G) pictorial depiction of the various allelic combinations that are analysed in (A-F). Note: In (A-F), one-way ANOVA followed by Dunnett’s test was used to determine the statistical significance. Multiple range comparison was performed, and the corresponding P-values are displayed in all the graphs (significant ones are in bold). Data obtained from field grown plants was used for the analysis in (A-F). The image (G) is partly created using biorender (https://biorender.com/). *bh^t^-A1+BH^t^-A3+GPC-B1*: Recombinants with one locus for spike-branching and two loci for accelerated senescence; *bh^t^-A1+bh^t^-A3+gpc-B1*: Recombinants with two loci each for spike-branching and delayed senescence. *Bh^t^-A1+bh^t^-A3+gpc-B1*: Recombinants without any spike-branching alleles, but contain two loci for delayed senescence.

**Fig. 8.**
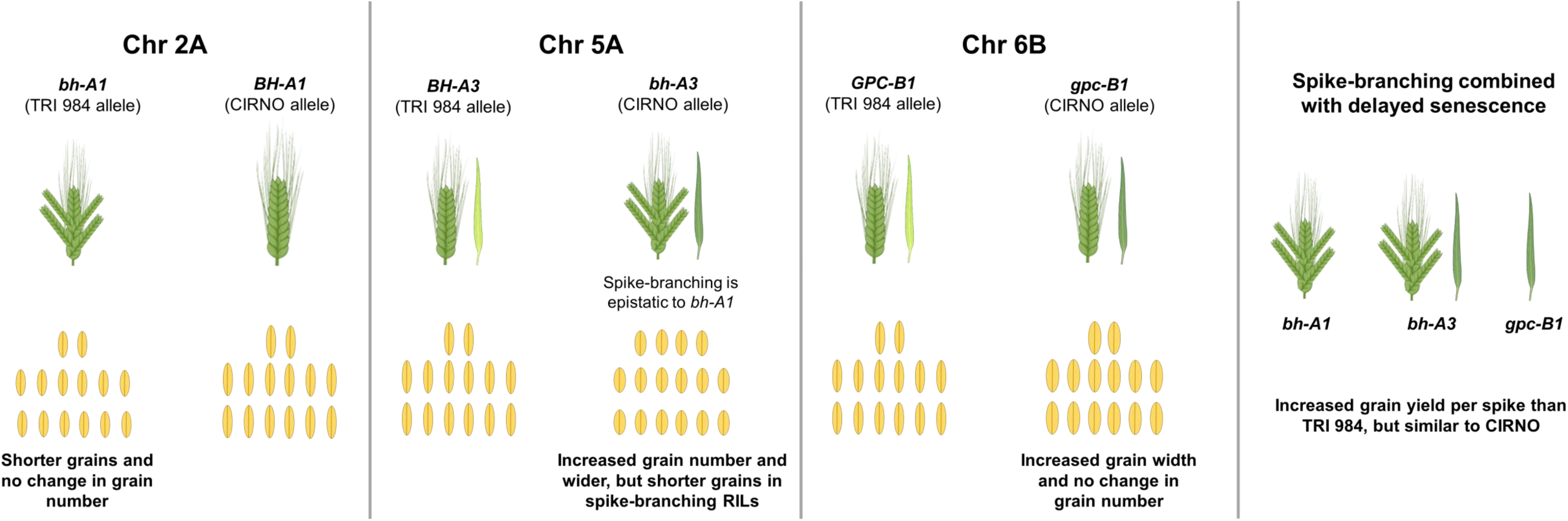
Summary of the physiological and genetic basis of interaction among *bh^t^-A1*, *bh^t^-A3*, and *gpc-B1* in the current landrace-elite recombinant population. Note: The image is partly created using biorender (https://biorender.com/).

## DISCUSSION

Over the course of domestication and breeding, grain yield determinants such as grain number and grain weight, but also grain quality traits under both favourable and stressful conditions, were the primary selection targets in all major cereal crops, including wheat (McSteen and Kellogg, 2022; Voss-Fels *et al*., 2019). For instance, the selection of the semi-dwarf *Rht-1* allele was a vital driver of the ‘green revolution’ in wheat (Peng *et al*., 1999); likewise, the prevalence of the less functional *GNI-A1* allele enabled higher floret fertility in the modern wheat cultivars (Golan *et al*., 2019; Sakuma *et al*., 2019). However, substantial genetic yield gaps [the difference between the genetic yield potential of a crop in a particular environment to that of the potential yield of the current local cultivar] suggest the presence of untapped genetic diversity for enhancing wheat grain yield (Senapati *et al*., 2022). Grain yield can be optimised by fine-tuning various developmental processes (Mathan *et al*., 2016) and introducing ‘drastic variations’ in crop breeding (Abbai *et al*., 2020). The genetic pathways that coordinate inflorescence architecture are dissected in staple grasses (Kellogg, 2022; Koppolu *et al*., 2022), which might have relevance for minimising the genetic yield gap.

Here, we considered the case of spike-branching Miracle-Wheat as a potential option for increasing sink strength (more spikelets and grains per spike). However, the genetic analysis of the TRI 984 x CIRNO recombinants revealed a couple of significant limitations. Firstly, we recorded inconsistencies in the expressivity (degree) of spike-branching (in the RILs that carried similar QTLs/alleles; Figs. 3B, 4B & S8B). Likewise, large variations in spike-branching intensity (Fig. S5H) and eventually grain number per spike (Fig. 1E) was also observed in TRI 984. (Wolde *et al*., 2021) reported that the expressivity of spike-branching in a particular genotype was higher in the outer rows as opposed to the inner rows of the plot. However, no new QTLs were mapped that specifically explained such differences. One explanation might be that field-grown plants experience competition for various resources, including light (Huber *et al*., 2021; Postma *et al*., 2021), especially in the inner rows (Rebetzke *et al*., 2014). Furthermore, we had two genotypes (three rows each) in 1.5 m^2^ plots in the current study and the neighbouring genotype were not the same in all the three replications, which might have affected spike-branching differently due to asymmetric plant-plant competition. In this regard, future studies investigating the response of various source and sink component traits in high-density monoculture plots or simulated canopy shade (Golan *et al*., 2022) are required to uncover the genetic framework of plant-plant competition and its effect on spike-branching expressivity.

Next, (Poursarebani *et al*., 2015) reported that the *bh^t^-A1* locus increases grain number, but with a grain weight trade-off. Likewise, we also observed considerably smaller grains in the spike-branching genotypes (Figs. 1F&3D). However, in the current study, the spike-branching phenotype induced by *bh^t^-A1* had no effect on the final grain number (Figs. 1E&3C). Interestingly, there was no thousand-grain weight trade-off in the spike-branching Bellaroi x TRI 19165 semi-dwarf RILs (Wolde *et al*., 2021) and also in the Floradur NILs with supernumerary spikelets (Wolde *et al*., 2019); thus, warranting the analysis of source-sink dynamics in the non-canonical spike forms. Here, it is vital to emphasise the relevance of the post-anthesis (yield realisation) events, chiefly related to the transfer of assimilates to the previously established sink organs during grain filling (Murchie *et al*., 2023; Slafer *et al*., 2023). In this context, the senescence rate might have an impact on grain filling duration (Chapman *et al*., 2021b; Christopher *et al*., 2016; Hassan *et al*., 2021; Kichey *et al*., 2007; Li *et al*., 2022), i.e., extended photosynthesis leading to more assimilate production and allocation to the developing grains. But, final grain weight was not strongly related with starch/sugar levels or the corresponding enzymatic capacity in 54 diverse wheat genotypes, but it might be a function of early developmental events (Fahy *et al*., 2018). In the current study, we found that higher grain number per spikelet (Fig. 2C) and grain weight (Fig. 2D) is associated with delayed flag leaf, peduncle and spike senescence. As expected, the observed effect of senescence rate might be because of the differences in various sink strength-related traits such as spikelet number per spike (spike-branching) (Fig. 3B), and floret number per spikelet (Fig. S9B) in our RIL population. As the sink strength increased, perhaps the extended photosynthetic period was meaningful for influencing the final grain yield. This trend further establishes the rationale for understanding the genetic and molecular framework of source and sink-related component traits to enable grain yield gains (Brinton and Uauy, 2019; Reynolds *et al*., 2022). With this, the favourable alleles explaining the source-sink dynamics might assist in improving grain number and grain weight in the spike-branching genotypes. Here, we analysed the interactions among *bh^t^-A1*, *bh^t^-A3,* and *gpc-B1*; the *bh^t^-A1* and *bh^t^-A3* loci regulated spike-branching, but also source strength, while *gpc-B1* delayed senescence rate and increased thousand-grain weight (Figs. 6&7). Transcriptional analysis of WT and *NAM* (*GPC*) RNAi lines revealed differential regulation of genes related to various processes, including photosynthesis and nitrogen metabolism, during flag leaf senescence (Andleeb *et al*., 2022). Our preliminary genetic evidence indicates that *gpc-B1* might function independently of the spike-branching associated loci (Fig. S13A&B). In any case, as speculated, the spike-branching RILs with an extended photosynthetic period (delayed senescence) had considerably higher grain yield (per meter row) as opposed to branched spike genotypes that senesced early (Fig. 6D). In this case, the stay-green spike-branching RILs had about 11 additional grains per spike (SEM: ±3.17) (Fig. 6A). However, we believe that the grain number difference might be due to the interaction between floret number and flag leaf senescence, which is mediated by the *bh^t^-A3* locus; the CIRNO allele increased florets per spikelet (Fig. S9B) and delayed flag leaf senescence (Fig. 4C). The pre-anthesis floret degeneration and post-anthesis flag leaf senescence might share a common genetic basis thereby primarily affecting the tip of the respective organs, i.e. spikelet meristem/rachilla and flag leaf, respectively. Therefore, it is conceivable that the underlying gene might have a pleiotropic effect on floret degeneration and flag leaf senescence, thus explaining the grain number difference. Recently, enrichment of senescence related transcripts has been reported during pre-anthesis tip degeneration in barley spikes (Shanmugaraj *et al*., 2023), which further supports our hypothesis. Then in our field experiments conducted at IPK-Gatersleben (2021 and 2022), we found an 8.5% (SEM: ±3.11%) increase in average grain weight (Fig. 6B) in the spike-branching genotypes that senesce late. The 2.53% (SEM: ±1.01%) rise in grain width (Fig. S12A) majorly contributed to the grain weight difference, as the grain length remained unaffected (Fig. S12B). Incidentally, it was found that grain width increased during wheat evolution under domestication (Gegas *et al*., 2010). Besides, it might be interesting to evaluate the effect of expansin genes in the spike-branching lines as the ectopic expression of *TaExpA6* increased grain length (Calderini *et al*., 2021).

Further, we would like to emphasise certain limitations in our experimental setup: we used relatively small plots (only 1.5 m^2^) with two genotypes in one plot; therefore, the influence of the border effect (Rebetzke *et al*., 2014) cannot be excluded in grain yield per row calculations and besides, the evaluated population are landrace-elite recombinants, that might create another bias in the observed yield increase. Although there is a significant increase in grain number per five spikes and grain weight in the stay-green spike-branching recombinants, the actual yield advantage might be better understood by evaluating the effect in isogenic backgrounds (NILs) and larger plots in multiple environments. In this context, we are developing spike-branching CIRNO NILs for these follow-up experiments. Another trade-off associated with extending the grain filling duration that is not addressed here is its likely impact on grain nutrition profile; the functional *NAM-B1* allele improves grain protein, iron and zinc content by accelerating the senescence process (Uauy *et al*., 2006). Then, the status of the stay-green spike-branching RILs under unfavourable conditions is also beyond the scope of the current study; however, previous reports indicate a positive effect of stay-green phenotypes on wheat grain yield under drought and heat (Lopes and Reynolds, 2012). Similarly, delay in senescence led to higher grain number and tiller number but lower thousand-grain weight under nitrogen-limiting conditions (Derkx *et al*., 2012). In addition, a recent simulation study indicates the advantage of cultivating late-maturing wheat varieties in future climate scenarios (Minoli *et al*., 2022), suggesting that a delay in senescence rate might eventually be beneficial.

## CONCLUSION

The physiological and genetic analysis of TRI 984xCIRNO recombinants revealed that i. extended verdant flag leaf, peduncle and spike led to higher grain yield per spike as the traits influencing sink strength segregated, including spike-branching; ii. we identified three QTL regions–on Chr 2A (*bh^t^-A1*), Chr 5A (*bh^t^-A3*) and Chr 6B (*gpc-B1*) that were associated with source-sink strength in the current bi-parental population; iii. upon analyzing their various allele combinations, it was found that an increase in grain number and grain weight is predominantly possible among the stay-green, spike-branching genotypes. iv. Finally, as wheat grain yield is also sink-limited, we propose that introducing spike-branching as a breeding target might enable advancing genetic gains while minimising the gap between genetic yield potential and the actual realised yield. Although we provide insights into the genetic basis of grain yield determination in ‘Miracle-Wheat’, it is still necessary to understand the basis of inconsistencies in the degree of spike-branching within the same genotype but also in diverse genetic backgrounds. To achieve this, tracking the source-strength dynamics during the early developmental stages might be necessary.

## ABBREVIATIONS

Chr: Chromosome
RILs: Recombinant inbred lines
QTL: Quantitative trait locus

## ACKNOWLEDGEMENT

We are grateful to Franziska Backhaus, Corinna Trautewig, Kerstin Wolf, Sonja Allner, Ellen Weiss, Ingrid Marscheider, and Angelika Püschel for their excellent technical assistance; Roop Kamal for helping with the field design; Prof. Nils Stein for his valuable comments on the project; Dr. Gemma Molero for sharing CIRNO grains; all members of Plant Architecture research group for the fruitful discussions and also for the support during harvest; Peter Schreiber and the team of gardeners for managing the field operations.

## AUTHOR CONTRIBUTIONS

TS acquired funding and supervised the project. RA continued to develop the population further, generated the data, analysed and interpreted the results. TS and GG guided in analysis and interpretation of the results. FHL conducted the field experiment at the University of Hohenheim. RA wrote the manuscript with inputs from all the co-authors.

## CONFLICT OF INTEREST

The authors declare that there is no conflict of interest.

## FUNDING

TS thanks the European Fund for Regional Development (EFRE), the State of Saxony-Anhalt within the ALIVE project, grant no. ZS/2018/09/94616, the HEISENBERG Program of the German Research Foundation (DFG), grant no. SCHN 768/15-1 and the IPK core budget for supporting this study. GG was funded through the Alexander von Humboldt Foundation postdoctoral fellowship program.

## DATA AVAILABILITY

Dataset S1 (48 excel sheets arranged figure-wise) and Dataset S2 (43 excel sheets arranged figure-wise) contain the data used for testing statistical significance for all the corresponding main and supplementary figures respectively.

## Notes

### Competing Interest Statement

The authors have declared no competing interest.

### Summary of Updates

Fig.7 is newly added to compare the spike grain yield data of the favourable RILs with the parents.

