## Supplemental Figures (S1-S13) for "Balancing grain yield trade-offs in ‘Miracle-Wheat’"

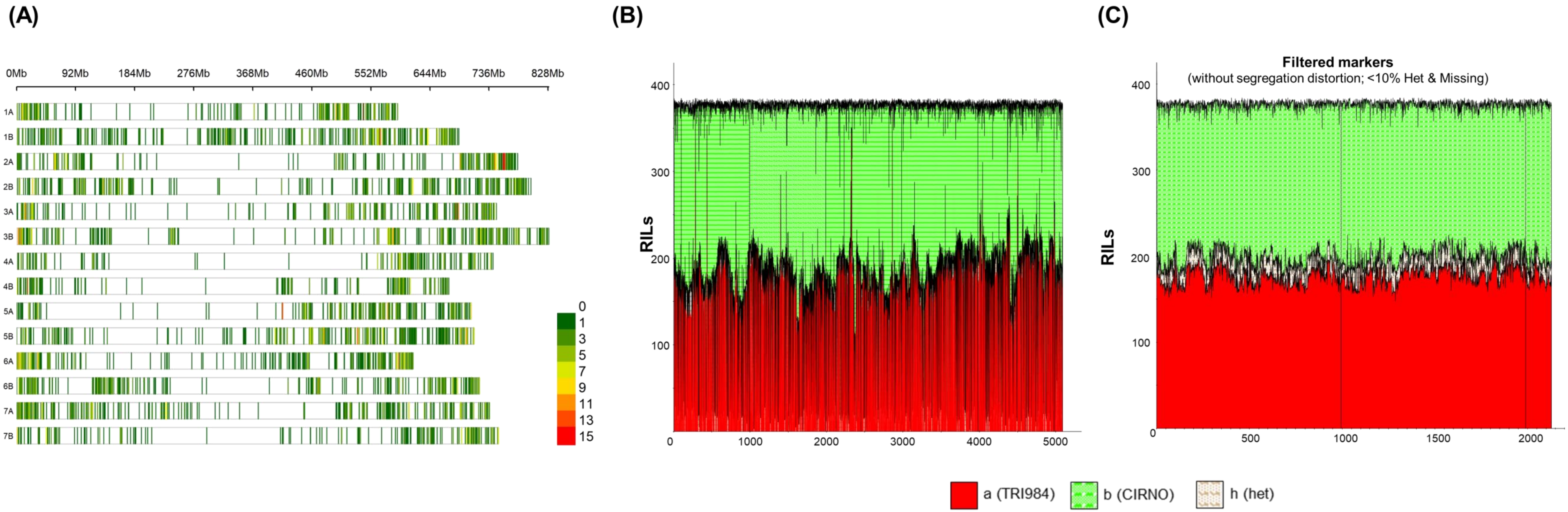


**Fig. S1.** The RIL population was genotyped using the 25K array. (A) a total of 5,089 polymorphic markers were found across the genome, (B) but with segregation distortion and (C) eventually, a filtered set of 2,089 markers was developed.

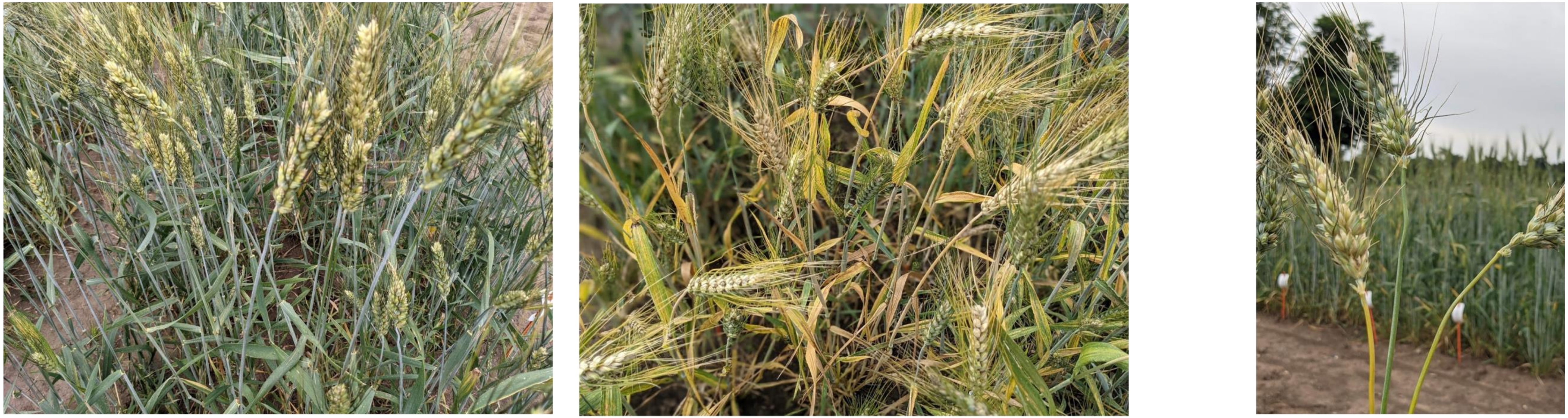

**(A)**

Flag leaf senescence Score '1'
(HD+30D)

Flag leaf senescence Score '4'
(HD+30D)

**(B)**

**Fig. S2.** Tracking the progression of senescence rate at 30 days after heading. (A) Phenotypes of flag leaves with delayed and accelerated senescence. (B) A gradient of peduncle yellowness was observed; however, the population was classified only into two categories in the current study.

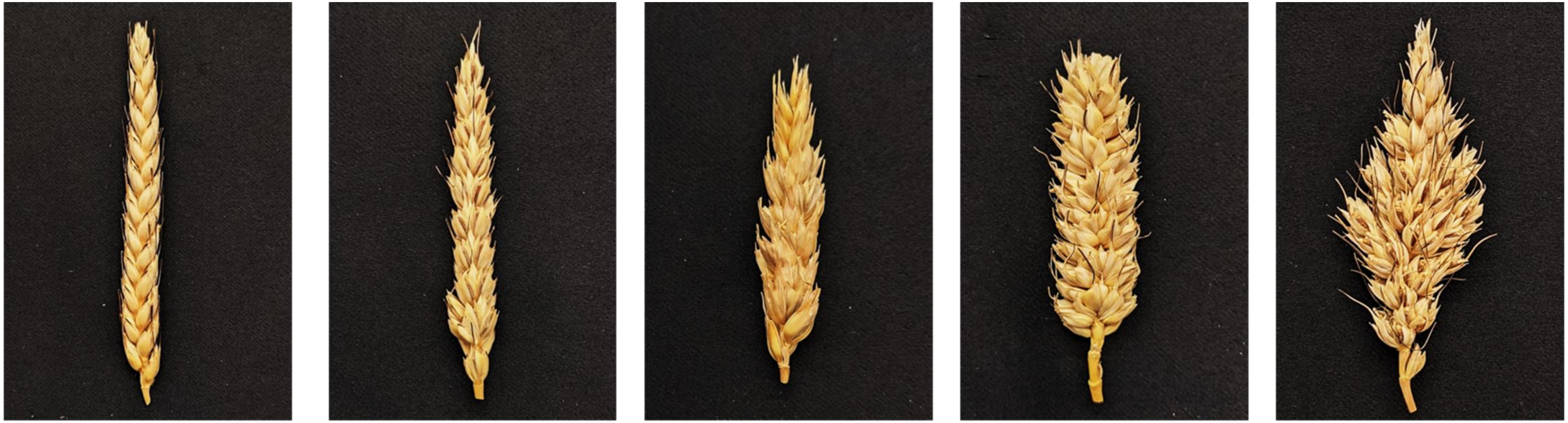

0

Standard spike with no supernumerary spikelets

1

Supernumerary spikelets only in the basal part of the spike

2

Supernumerary spikelets in the central and basal part of the spike

3

Supernumerary spikelets throughout the spike

4

True spike-branching phenotype

**Fig. S3.** Dosage-based scoring method (0-4 point scale) for spikes with supernumerary spikelets and spike-branching phenotypes.

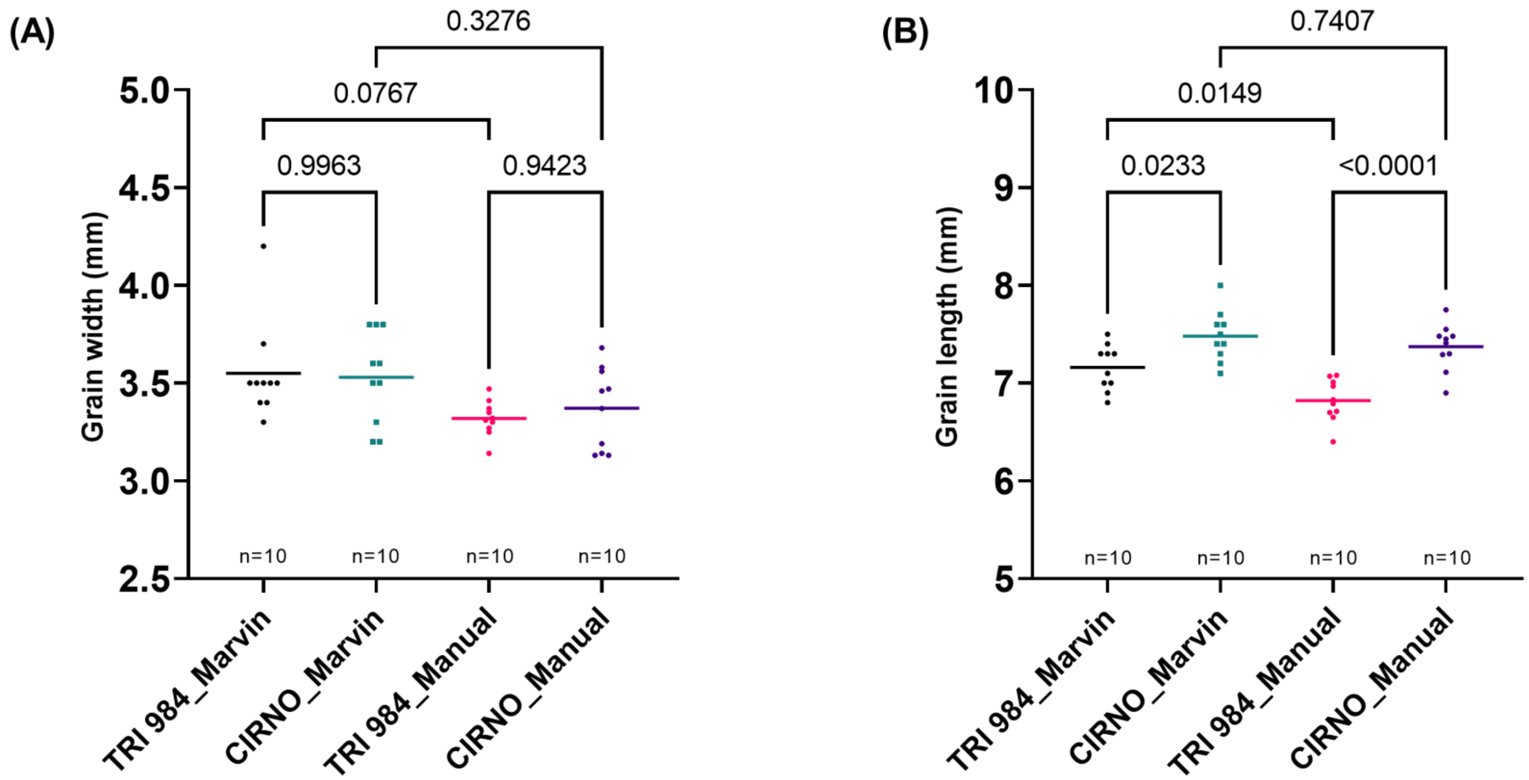


**Fig. S4.** Validation of Marvin results. (A) The grain width and (B) grain length results obtained from the Marvin grain analyser were confirmed manually using a Vernier caliper. Although there was a difference in absolute values, the trend between the parental lines remained the same; thus, it might not affect the QTL mapping analysis. Note: Data obtained from field grown plants was used for the analysis in (A-B).

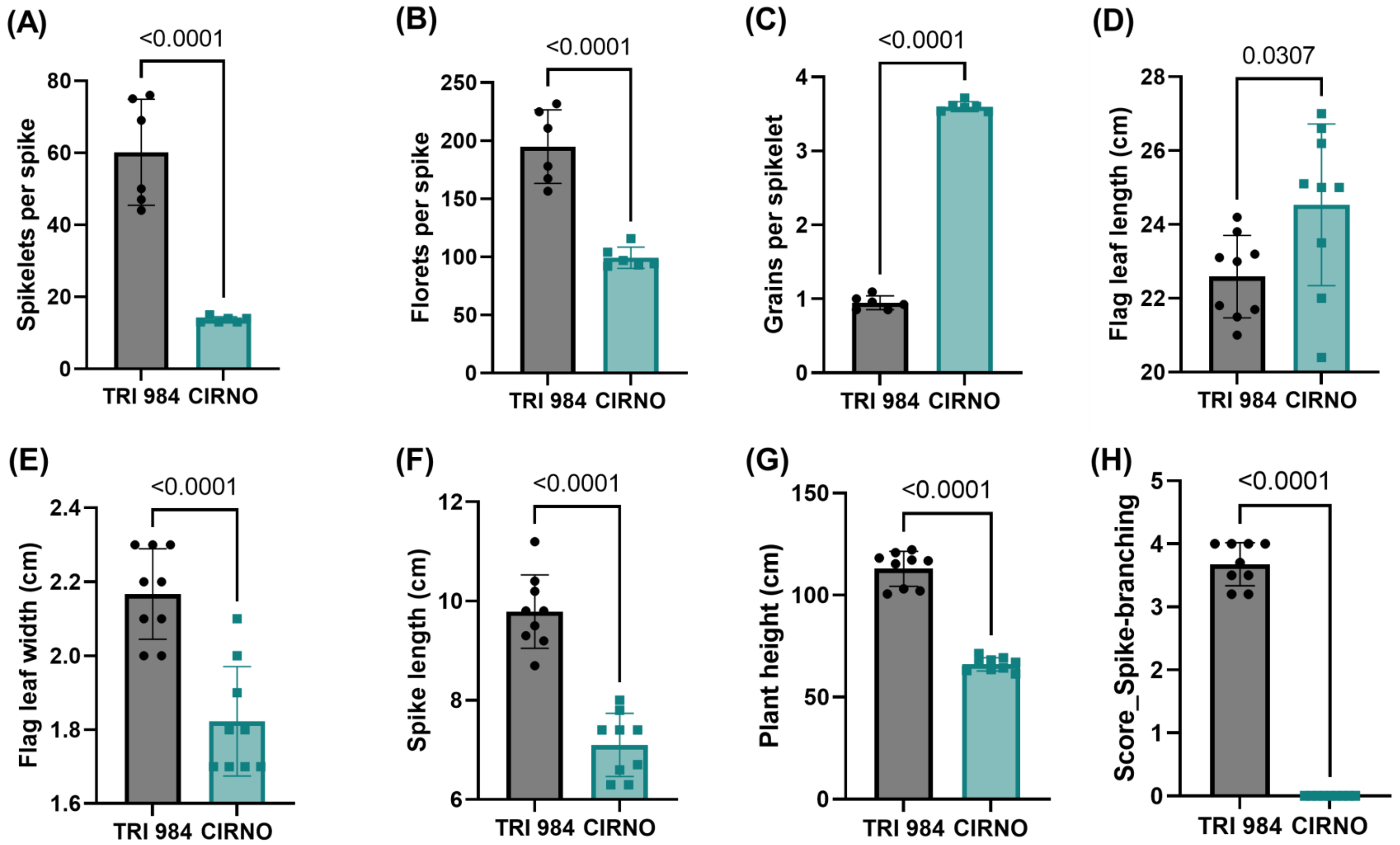


**Fig. S5.** TRI 984 vs CIRNO. (A) Spikelets per spike, (B) Florets per spike, (C) Grains per spikelet, (D) Flag leaf length, (E) Flag leaf width, (F) Spike length, (G) Plant height, and (H) Spike-branching score. Note: Data obtained from field grown plants was used for the analysis in (A-H).

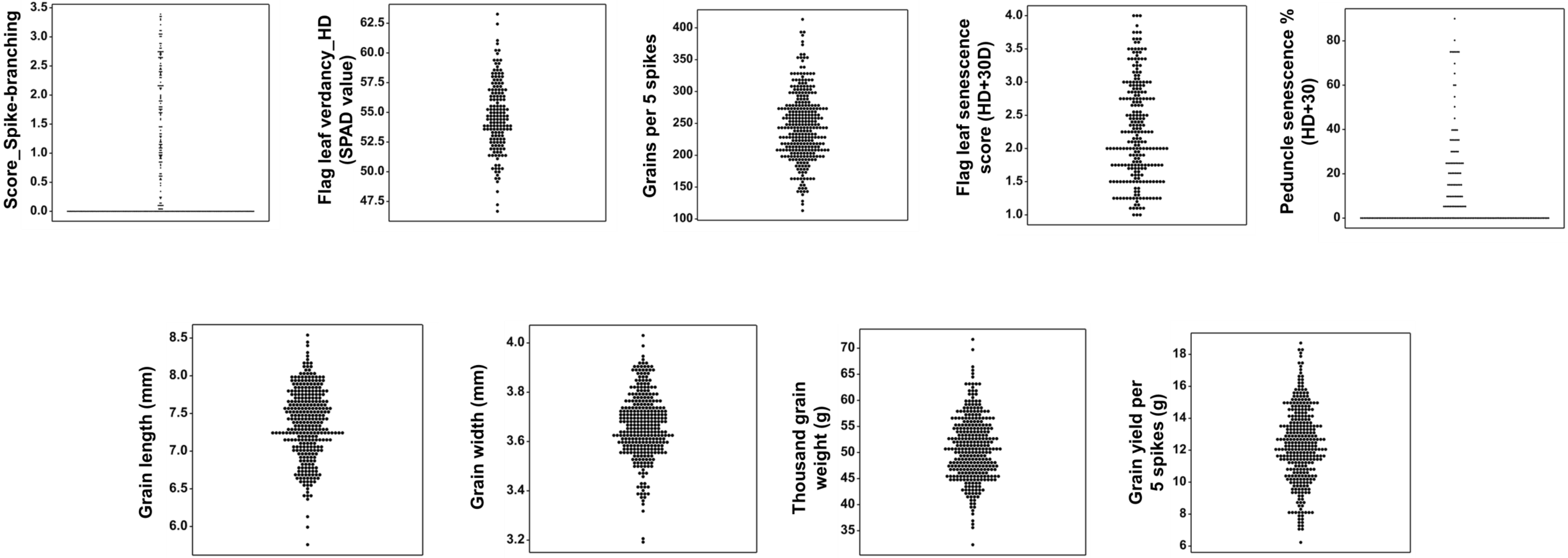


**Fig. S6.** Phenotypic distribution of various plant and spike architectural traits across the RIL population. Note: Data obtained from field grown plants was used for the analysis.

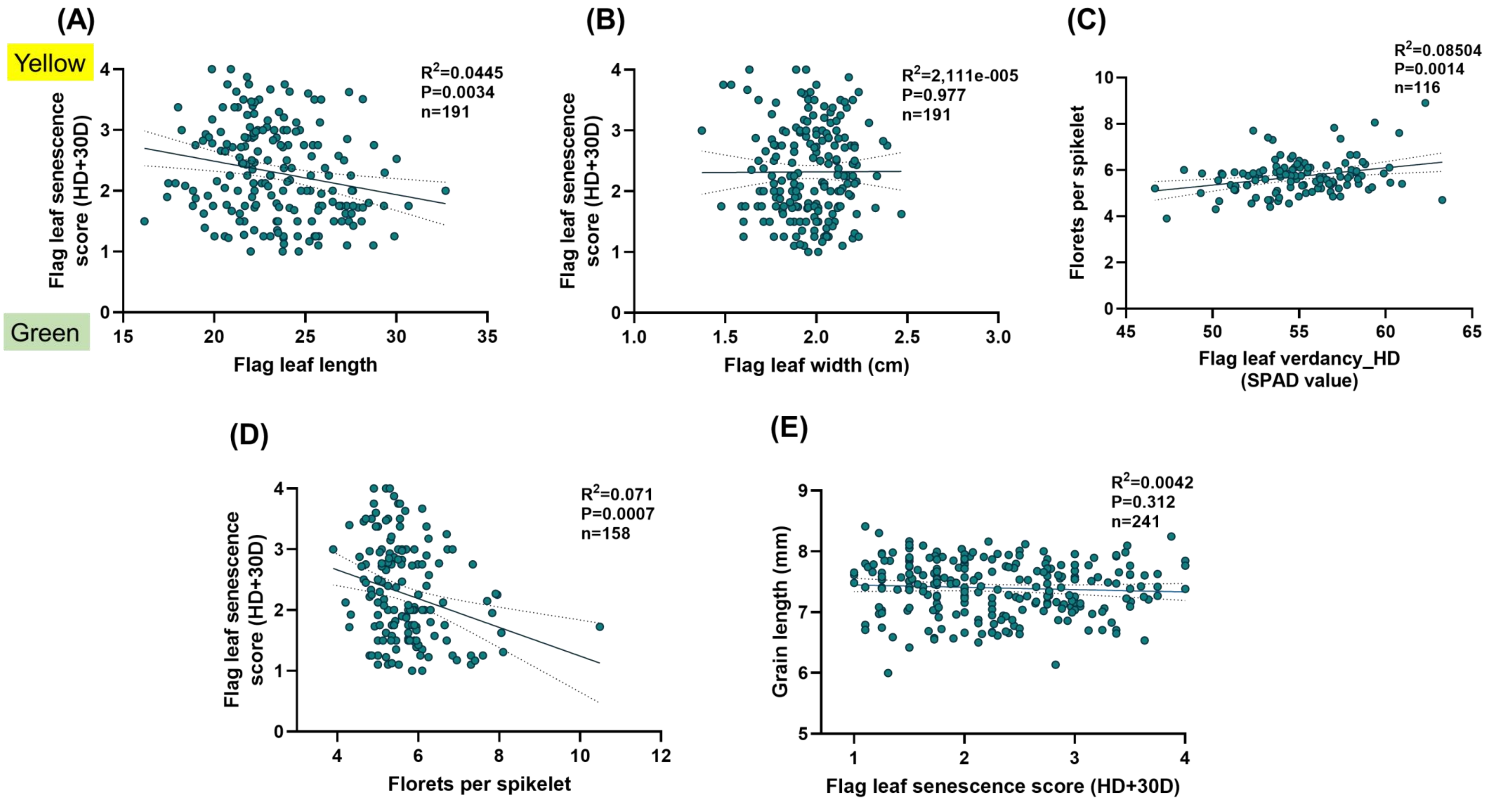


**Fig. S7.** Relationship between various traits across the population. (A) Flag leaf length vs Flag leaf senescence, (B) Flag leaf width vs Flag leaf senescence, (C) Flag leaf verdancy at heading vs Florets per spikelet, (D) Florets per spikelet vs Flag leaf senescence, and (E) Flag leaf senescence vs Grain length. Note: Data obtained from field grown plants was used for the analysis in (A-E).

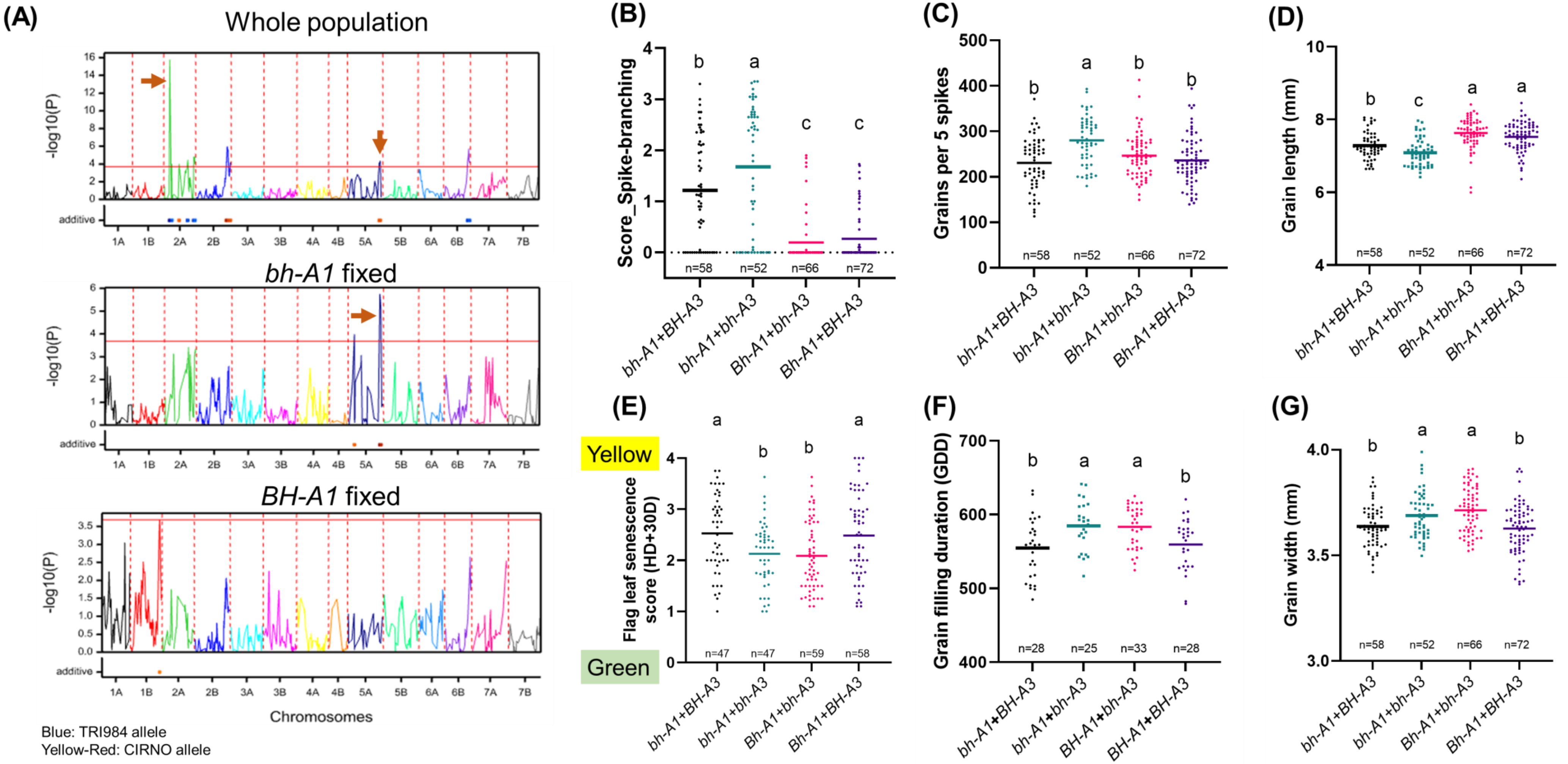


**Fig. S8.** Spike-branching phenotype explained by *bh-A3* is epistatic with *bh-A1*. (A) QTL mapping using the whole population, but also with two subsets developed by fixing *bh-A1* and *BH-A1*. (B) The spike-branching expressivity, (C) grains per 5 spikes and (D) Grain length in various allele combinations of the two QTLs. However, the role of *bh-A3* in affecting (E) flag leaf senescence, (F) Grain filling duration and (G) Grain width is independent of *bh-A1*. Note: Data obtained from field grown plants was used for the analysis in (A-G).

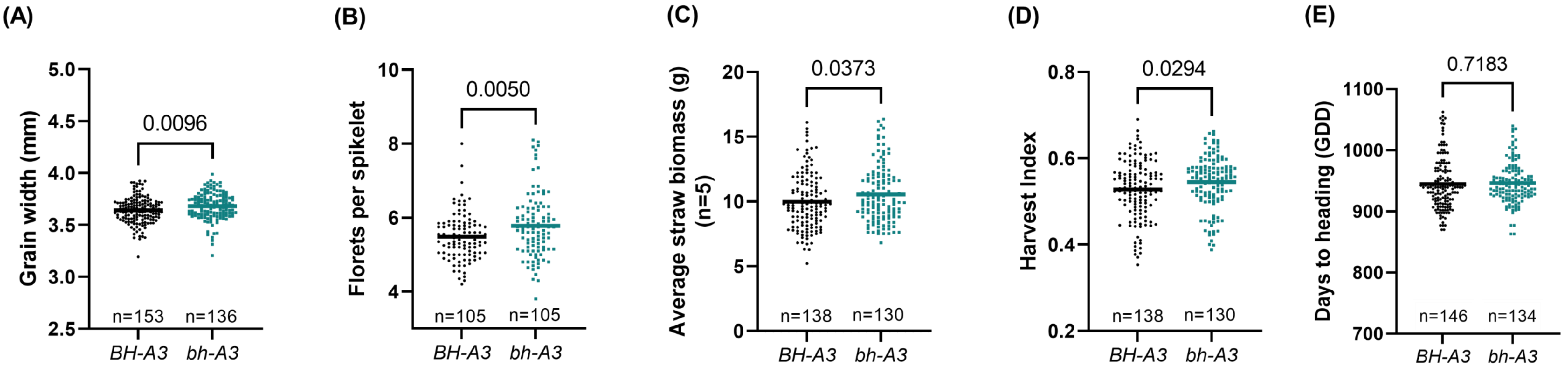


**Fig. S9.** *bh-A3* influences (A) Grain width, (B) Florets per spikelet, (C) Straw biomass, (D) Harvest index and (E) has no effect on days to heading. Note: Data obtained from field grown plants was used for the analysis in (A-E).

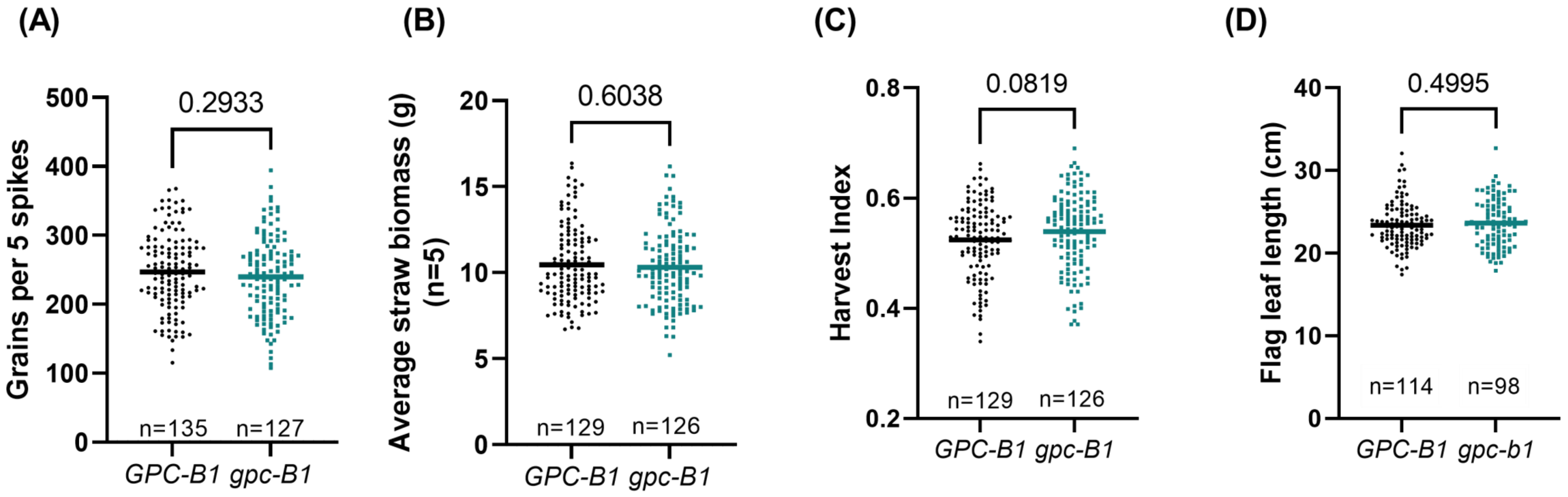


**Fig. S10.** *gpc-B1* does not influence (A) Grains per 5 spikes, (B) Straw biomass, (C) Harvest index and (D) Flag leaf length. Note: Data obtained from field grown plants was used for the analysis in (A-D).

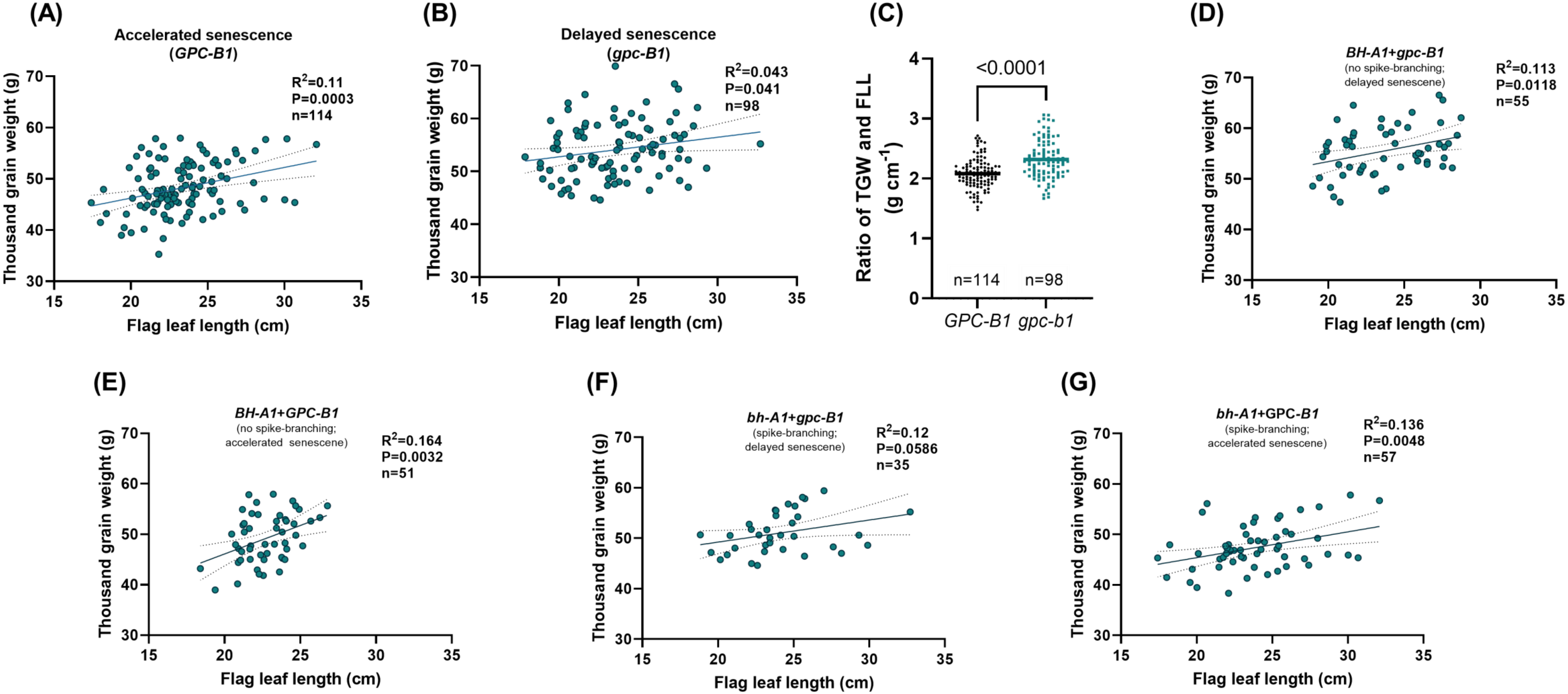


**Fig. S11.** Effect of flag leaf length and thousand grain weight in RILs carrying (A) *GPC-B1* and (B) *gpc-B1*. The contribution to thousand grain weight per unit length of flag leaves was higher in (C) *gpc-B1* compared to *GPC-B1*. (D-G) Various allele combinations of *gpc-B1* and *bh-A1* and their implications on the relationship between flag leaf length and thousand grain weight. TGW: Thousand grain weight and FLL: Flag leaf length. Note: Data obtained from field grown plants was used for the analysis in (A-G).

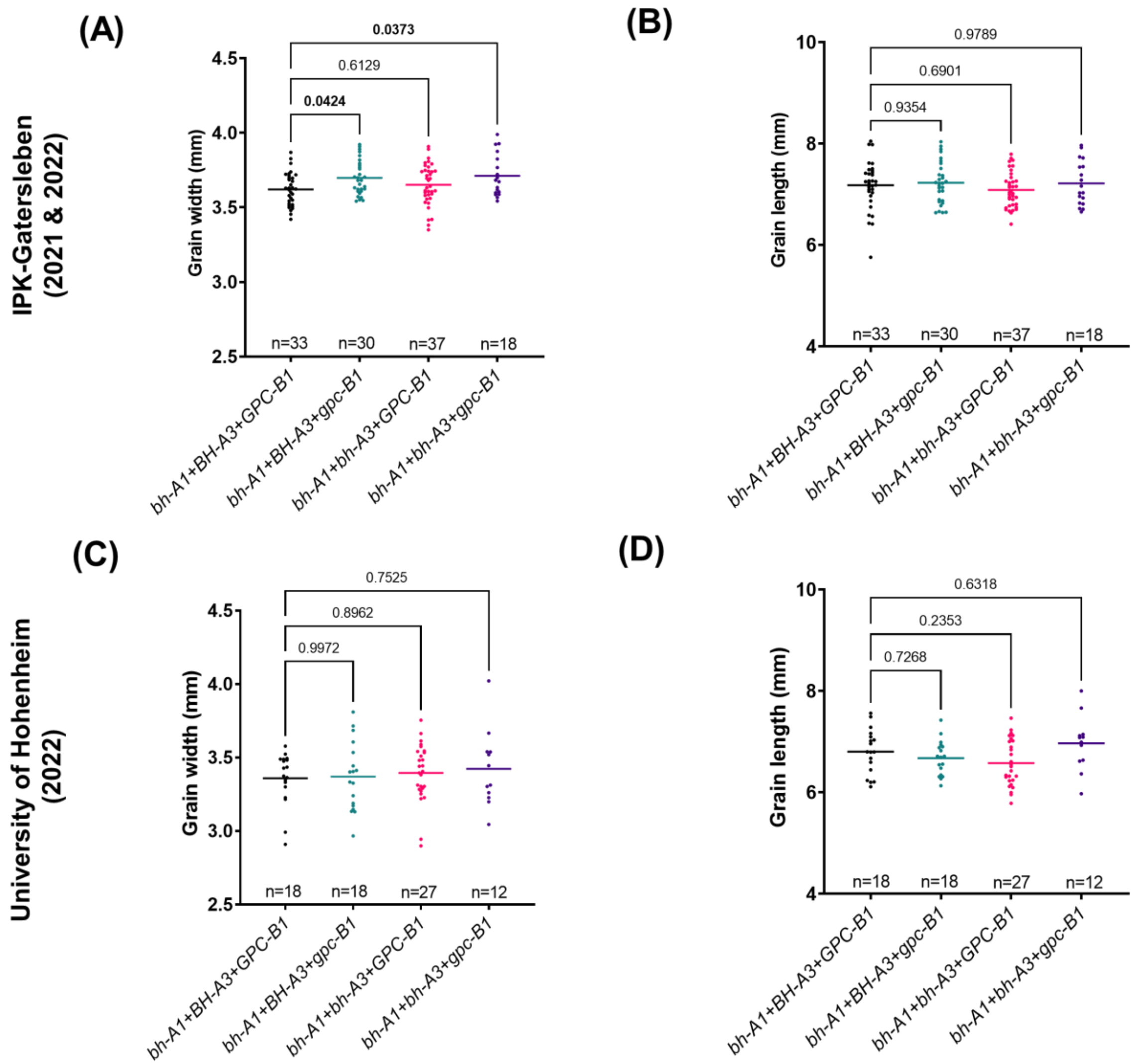


**Fig. S12.** Genetic interaction among *bh-A1*, *bh-A3* and *gpc-B1*. The effect of various allele combinations at IPK (A) Grain width, (B) Grain length and at University of Hohenheim (C) Grain width, (D) Grain length. Note: Data obtained from field grown plants was used for the analysis in (A-D).

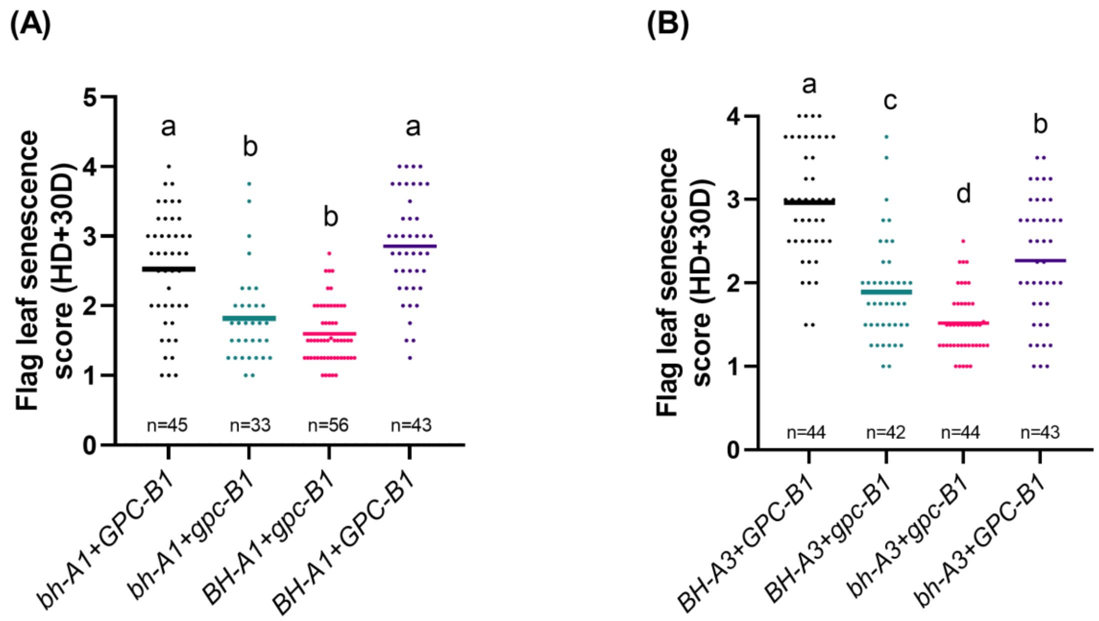


**Fig. S13.** The flag leaf senescence rate of various allele combinations revealed that (A) *gpc-B1* is independent of *bh-A1* and (B) *bh-A3* and *gpc-B1* exhibit additive effect. Note: Data obtained from field grown plants was used for the analysis in (A-B).
